# Niche engineering drives early passage through an immune bottleneck in progression to colorectal cancer

**DOI:** 10.1101/623959

**Authors:** Chandler D. Gatenbee, Ann-Marie Baker, Ryan O. Schenck, Margarida P. Neves, Sara Yakub Hasan, Pierre Martinez, William CH Cross, Marnix Jansen, Manuel Rodriguez-Justo, Andrea Sottoriva, Simon Leedham, Mark Robertson-Tessi, Trevor A. Graham, Alexander R.A. Anderson

## Abstract

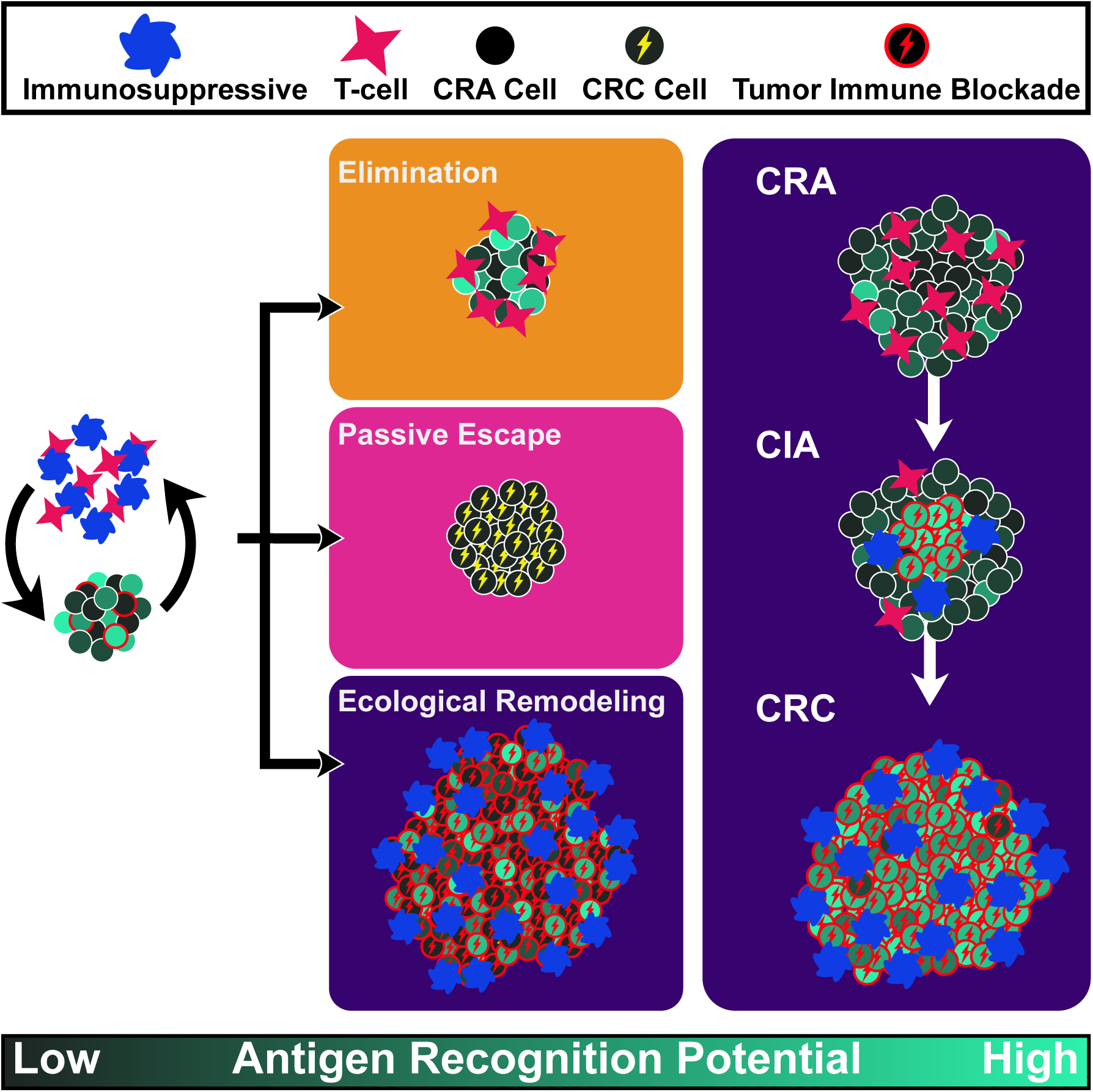

Colorectal cancer develops from its precursor lesion, the adenoma. The immune system is hypothesized to be key in modulating progression, but tumor-immune eco-evolutionary dynamics remain uncharacterized. Here, we demonstrate a key role for immune evasion in the progression of human benign disease to colorectal cancer. We constructed a mathematical model of tumor-immune eco-evolutionary dynamics that predicted ecological succession, from an “immune-hot” adenoma immune ecology rich in T cells to an “immune-cold” carcinoma ecology, deficient in T cells and rich in immunosuppressive cells. Using a cross-sectional cohort of adenomas and carcinomas, we validated this prediction by direct measurement of the tumor-immune ecology using whole-slide 10-marker immunohistochemistry (IHC), and analysis of neoantigen clonal architecture multi-region exome sequencing data. Changes in immune ecology relax selection against antigens with high recognition potentials. This study indicates that immune surveillance represents a key evolutionary bottleneck in the evolution of colon cancer.

## Introduction

The classical model of colorectal carcinogenesis is the adenoma-carcinoma pathway that describes the accumulation of (epi)mutations in benign (non-invasive) adenomas which underpin the development of invasive carcinoma (Muto, Bussey, & Morson, 1975). However, while the risk of developing colorectal cancer (CRC) is certainly increased by adenoma formation (Carvajal-Carmona et al., 2013), it appears that few adenomas actually progress to cancer in a human lifetime. Bowel cancer screening programs detect approximately five ‘high risk’ adenomas for every cancer found (Logan et al., 2012; Zauber et al., 2012), and longitudinal endoscopic surveillance of adenomas reveals that less than 2% of adenomas progress to cancer within three years (Hofstad et al., 1996). Consequently, there appears to be a substantial ‘evolutionary hurdle’ that must be overcome for an adenoma to become invasive.

Immune predation is known to modulate and indeed suppress neoplastic growth (Grivennikov, Greten, & Karin, 2010). The evolution of immune evasion is therefore likely to be a key barrier on the evolutionary path to malignancy. Newly arising somatic mutations in a tumor have the potential to generate neoantigens, which can then serve as targets for recognition and destruction by the cells of the immune system (in particular CD8+ cytotoxic T lymphocytes). However, due to the selection pressure imposed by the immune response, tumor cells often evolve strategies to avoid elimination in a process known as immunoediting (Dunn, Bruce, Ikeda, Old, & Schreiber, 2002; Grasso et al., 2018). Such immune evasion has been described as a hallmark of cancer (Hanahan & Weinberg, 2011), and there are many mechanisms by which tumor cells may escape immune predation, including, but not limited to, suppression of the cytotoxic T-cell response via expression of programmed death-ligand 1 (PD-L1), recruitment of immunosuppressive cells such as macrophages and neutrophils, and disruption of the antigen presentation machinery (Coffelt, Wellenstein, & de Visser, 2016; Khong & Restifo, 2002; Schreiber, Old, & Smyth, 2011; Vinay et al., 2015).

In colorectal cancer, multiple lines of evidence suggest a critical role for immunological surveillance in regulating tumor growth. The degree of tumor-infiltrating T cells is highly prognostic in CRC, with greater infiltration associated with a better prognosis (Galon et al., 2006; Pages et al., 2009), and moreover non-metastatic CRCs have an increased level of T-cell infiltration as compared to metastatic CRC (Pages et al., 2005). Genomic analysis reveals that a higher predicted neoantigen burden is associated with increased tumor lymphocyte infiltration (Angelova et al., 2015; Giannakis et al., 2016). Microsatellite unstable cancers (MSI+), that have a particularly high density of infiltrating lymphocytes (Dolcetti et al., 1999) and generate a high number of neoepitopes (Angelova et al., 2015; Giannakis et al., 2016) have generally good prognosis compared to MSI-tumors, and also are very responsive to programmed death 1 (PD-1) immune checkpoint blockade (Le et al., 2015). Genetic evidence of immune evasion is very common in CRCs (Grasso et al., 2018). Immune modulation studies in mouse models of CRC also provide support for a critical regulatory role of the immune system in colorectal carcinogenesis (B. G. Kim et al., 2006; Ngiow et al., 2011; Yu, Steel, Zhang, Morris, & Waldmann, 2010).

The progression from adenoma to cancer likely requires the accumulation of multiple (epi)genetic mutations. While the overall single nucleotide alteration (SNA) burden appears comparable between adenomas and cancers, including SNAs for putative driver genes (Cross et al., 2018), the burden of somatic copy number alterations (CNAs) is much higher in cancers. Intriguingly, few mutations in CRC show tumor stage specificity: only *TP53* mutation and large-scale CNA are enriched in cancers (Cross et al., 2018) Moreover, analysis of the evolutionary dynamics of sub-clones within CRC indicates a lack of subclonal selection (Williams et al., 2018) suggesting that at the point of invasion, the founder cancer cell had already acquired all the alterations necessary for its malignant phenotype and also that the bulk of tumor cells were not experiencing differential immune predation. Thus, malignant potential and immune evasion may be established together when cancer growth is initiated, and conceivably immune evasion could be the key phenotypic trait governing transition from adenoma to cancer.

Here, we have investigated the role of immune escape in the evolution of colorectal cancer from precursor adenoma lesions. We hypothesized that immune surveillance represents a key hurdle that prevents the development of invasive cells within a benign adenoma. The acquisition of mutations responsible for progression must be balanced against the risk of accumulating multiple neoantigens that lead to immune-mediated suppression of tumor growth. To investigate this idea, we developed a computational model that simulates tumor evolution under immune predation in order to determine the expected patterns of immune activity and intratumor antigenic heterogeneity (aITH) throughout tumor progression from benign to malignant. We then looked for these signatures in a cross-sectional cohort of colorectal adenomas as well as early and later stage cancers, using 10-marker immunohistochemistry (IHC) to characterize changes in the tumor ecology and spatial interactions, and exome sequencing to measure aITH. Our data point to a key role for immune evasion at the onset of malignancy.

## Results

### Ecological succession shapes antigenic intra-tumor heterogeneity

#### Modeling tumor-immune eco-evolutionary dynamics

Evolution is mathematically described by a branching process, and tumor-immune interactions can be understood as ecological relationships. Thus, we simulate these eco-evolutionary dynamics by constructing a stochastic branching process model inspired by Lotka-Volterra equations of competition, predation, and mutualism (Figure 1), wherein tumor cells are “prey”, T cells the “predator”, and immunosuppressive cells the “mutualists”. Clones, defined as the collection of cells that share the same mutations (including neoantigens), are the unit of selection in this model. With small probability, mutants are created during cell division, giving rise to new clones that inherit all parental mutations and carry an additional unique mutation associated with a neoantigen. Mutations potentially incur an evolutionary cost to the cell, since each neoantigen that arises from mutation stimulates immune attack (Coulie, Van den Eynde, van der Bruggen, & Boon, 2014). The strength of attack is determined by a randomly assigned recognition potential between 0 and 1 (uniformly distributed), where the higher the value the greater the rate of immune kill. While most mutations are deleterious through this neoantigen acquisition, we also model three types of possible beneficial somatically-acquired alterations: driver mutations, ability to block immune attack, and ability to recruit immunosuppressive cells, each of which is described in more detail below.

**Figure 1.**
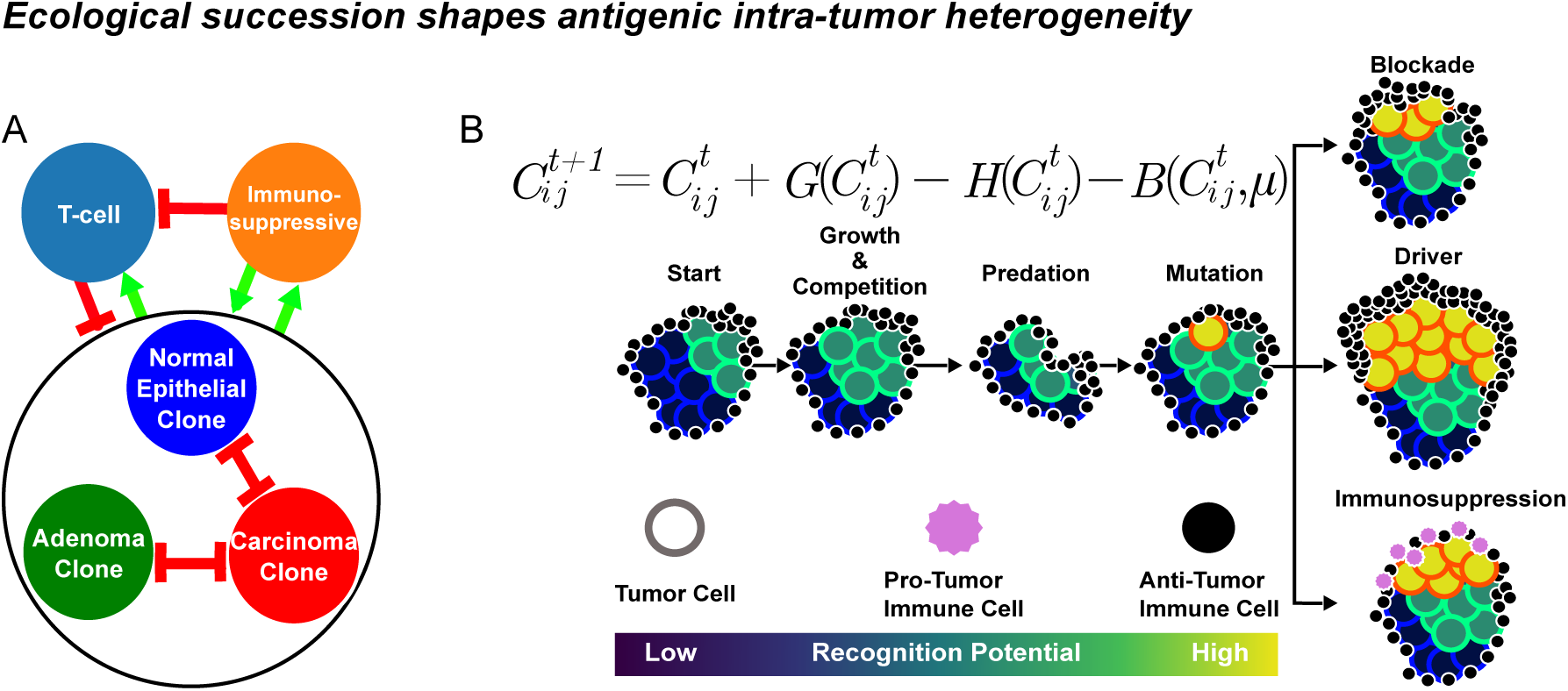
Tumor-immune eco-evolutionary dynamics are simulated using a branching process model inspired by Lotka-Volterra equations of competition, predation, and mutualism. **A** Cellular interactions. Red bars indicate inhibitory interactions and green arrows represent positive interactions. **B** Sequence of events in each time step of the model, where *C*_*ij*_ are antigenic clones of the three possible cellular “species” (subscript *i* for epithelial, adenoma, carcinoma), each with different antigenic clones *j*. *G*() is the function that governs clonal growth and competition, *H*() defines how many cells are lost to immune predation, and *B*() determines how many mutants are created by the clonal population. The model is initiated with a large population of homogenous non-immunogenic epithelial cells. Mutation may occur during division, resulting in the creation of a new clone in *i* which inherits all ancestral antigens and mutations from the parental clone. Each mutation is accompanied by the generation of a neoantigen, which stimulates additional attack by predatory T cells at a rate proportional to the neoantigen’s recognition potential assigned at the time of mutation. With low probability, three mutations may be acquired that are beneficial: 1) the ability to recruit immunosuppressive cells, which decrease T-cell attack while increasing growth rates; 2) the ability to block and thus reduce T-cell attack; and 3) acquisition of driver mutations, which increase division rates and carrying capacities. If an epithelial clone acquires two driver mutations it is considered an adenoma, which grows in a separate niche atop the epithelial tissue, limiting interactions between the two populations. Clones become carcinomas when they acquire four driver mutations. Carcinomas have the ability to grow both atop and into the epithelium, allowing carcinomas to invade and destroy epithelial and adenoma clones.

Clones belong to one of three “species”: normal epithelium (E), adenoma (CRA), and carcinoma (CRC), each with a specific division rate and carrying capacity (maximum population size) both of which increase through the progression to malignancy. We assume an epistatic model of progression (Fearon & Vogelstein, 1990), such that the acquisition of two driver mutations by an epithelial clone leads to an adenoma clone; acquiring a further two driver mutations creates a carcinoma clone. We also model competition between species that could occupy the same space in the colon: E and CRA clones have only weak interactions because adenomas grow superficially to the epithelium, whereas to account for tumor invasion we assume that CRC clones are strongly interacting with both E and CRA clones.

The collection of antigens carried by a clone determines its immune kill rate. Since all antigens are inherited, this means that, in the absence of an escape mechanism, all descendant clones experience the same or greater immune predation than any of their ancestors. We assume that the number of T cells per antigen is proportional to the number of cells that carry the antigen, scaled linearly by the antigen’s recognition potential. Thus, immune kill is determined using the clone’s size and the sum of the recognition potentials for each antigen, and not through explicit modeling of T-cell dynamics. A time lag is also included such that an immune response to the clone’s neoantigen is only activated after it becomes ‘detected’, where the probability of detection is determined by the total number of cells that carry the neoantigen and that neoantigen’s recognition potential (see Supplement for details). Once a neoantigen is detected, all clones that carry that same neoantigen experience increased predation.

A clone can *passively* escape immune predation by ‘getting lucky’ and only acquiring neoantigens with low recognition potentials. Alternatively, clones can *actively* mitigate immune attack via two mechanisms: immune blockade (e.g. PD-L1 or CTLA-4 expression), and/or suppression of the inflammatory immune response via recruitment of immunosuppressive cells, (e.g. M2 macrophages and neutrophils) (reviewed in (Coffelt et al., 2016; Khong & Restifo, 2002; Schreiber et al., 2011; Vinay et al., 2015)). Acquiring the ability to block immune attack has the effect of reducing the number of cells killed by immune predation. Recruitment of immunosuppressive cells can similarly reduce the kill rate, under the assumption that such suppressive cells reduce T-cell infiltration.

In addition to suppressing the anti-tumor immune response, immunosuppressive cells, such as macrophages and neutrophils, are also involved in wound repair, and are capable of remodeling the microenvironment to favor tumor growth (Coffelt et al., 2016; Galdiero, Varricchi, Loffredo, Mantovani, & Marone, 2018; J. Kim & Bae, 2016; Mantovani, 2014). Recruitment of these cells thus likely provides an evolutionary benefit to tumor cells by increasing both their growth rate and the maximum size of a clone. We assume that the density of immunosuppressive cells is locally increased on clones that have acquired the ability to recruit those cells. As such, the growth benefits are experienced only by clones capable of recruitment, and not the entire tumor.

To better understand how the strength of immune escape affects progression, we conducted a sweep over the parameters that control how much (i) immune blockade protects cells, (ii) immune suppression protects cells, and (iii) recruitment of immunosuppressive cells increases growth. Parameters were selected such that there was equal coverage of growth and combined protection (the effects of simultaneous blockade and suppression on protection are multiplicative). For each of these parameters, we ran the simulation until a carcinoma was sustained for one year. If this did not happen, the simulation was re-run, and the process repeated until a carcinoma existed for a full year. For each parameter set we thus have one simulation that generated a carcinoma that successfully evaded immune elimination, and potentially several simulations where an adenoma or carcinoma was eliminated by immune predation.

The dynamics of two representative simulations using different parameter sets can be found in Figure 2. Panel A shows a case where the tumor develops immunosuppression very early, allowing a broad range of recognition potentials to coexist. In contrast, the simulation in panel B does not allow for immunosuppression or blockade, forcing the tumor to ‘get lucky’ by only acquiring antigens with low recognition potential (note the scale of the immunogenicity axes). These examples highlight how aITH is affected by the ability to mitigate immune attack via niche engineering, as opposed to passive evasion through relying on low antigen burden.

**Figure 2.**
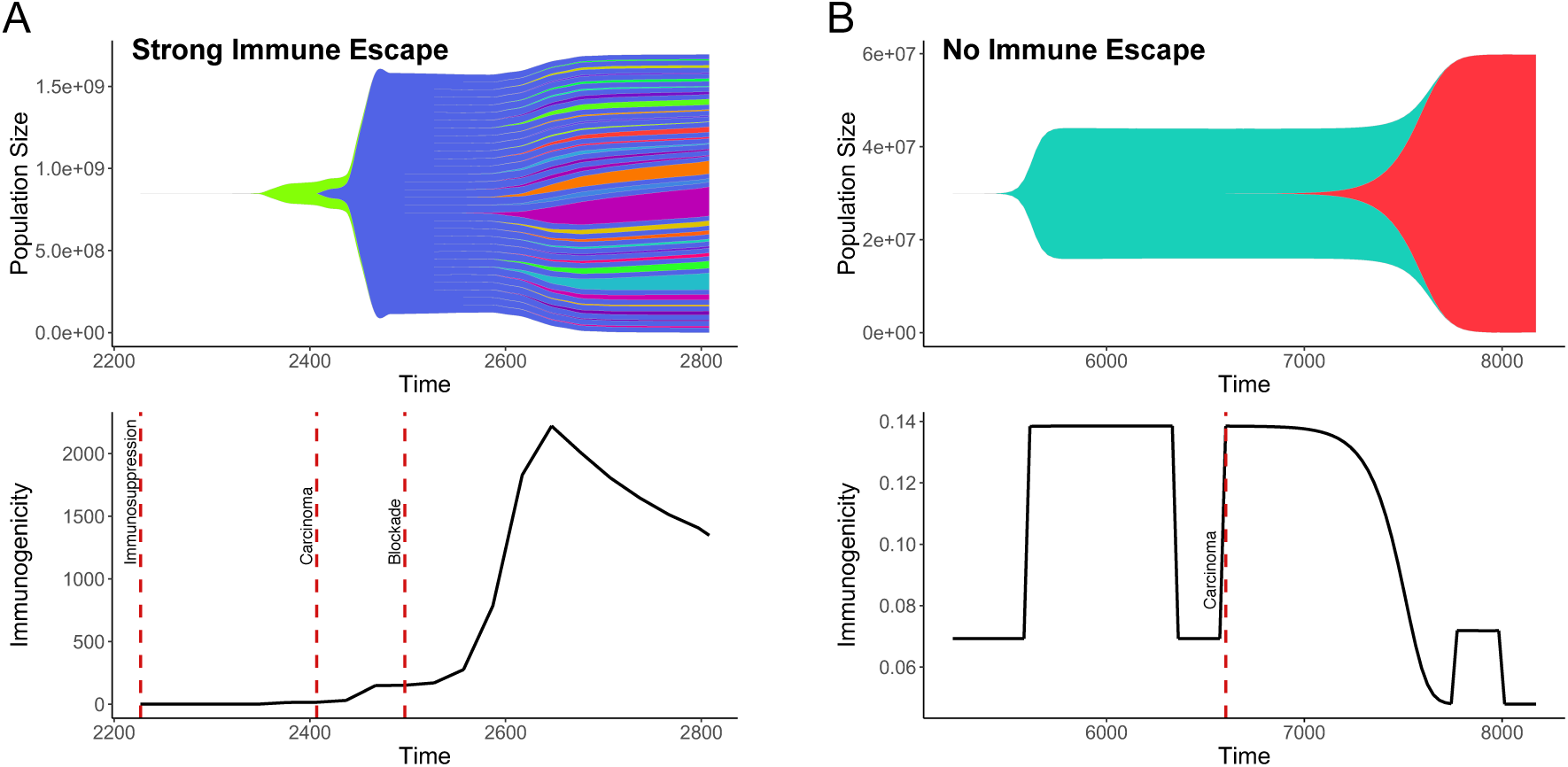
Simulated tumor-immune eco-evolutionary dynamics with (**A**) and without (**B**) immune escape. The top plot of each panel shows how aITH changes during tumor evolution. In these Muller plots, the height of each polygon represents the total number of cells that carry an antigen, which includes the source clone (i.e. the clone in which the antigen originated) and all of its descendants. The bottom plots of each panel show how the tumor’s immunogenicity, defined by as the product of antigen burden and the average recognition potential at any given time (note the three orders of magnitude difference in immunogenicity in the presence versus absence of immune escape). Dotted lines indicate important evolutionary events: the transition to carcinoma, the ability to recruit immunosuppressive cells, and T-cell attack blockade. *Remodeling of the immune ecology begins early and is essential for transformation*

By simulating tumor-immune eco-evolutionary dynamics, we characterized the immune ecology and the aITH that it shapes, before, during, and after escape from immune predation. We separately considered adenomas that were eliminated by immune predation (xCRAs), and those that progressed (CRAs). For each group, we examined the subset of adenomas that had not yet evolved the ability to mitigate immune attack. Selection against antigens with high recognition potentials (Figure 3A vertical axis) is intense. Only tumors that ‘get lucky’ by carrying antigens with low recognition potentials survive (CRA), while those carrying antigens with high recognition potentials are rapidly eliminated by strong immune predation (xCRA). Correspondingly, clonal diversity (Figure 3B) in adenomas that are eliminated (xCRAs) is uniformly low, whereas ‘successful’ CRAs could exhibit much greater levels of clonal diversity.

**Figure 3.**
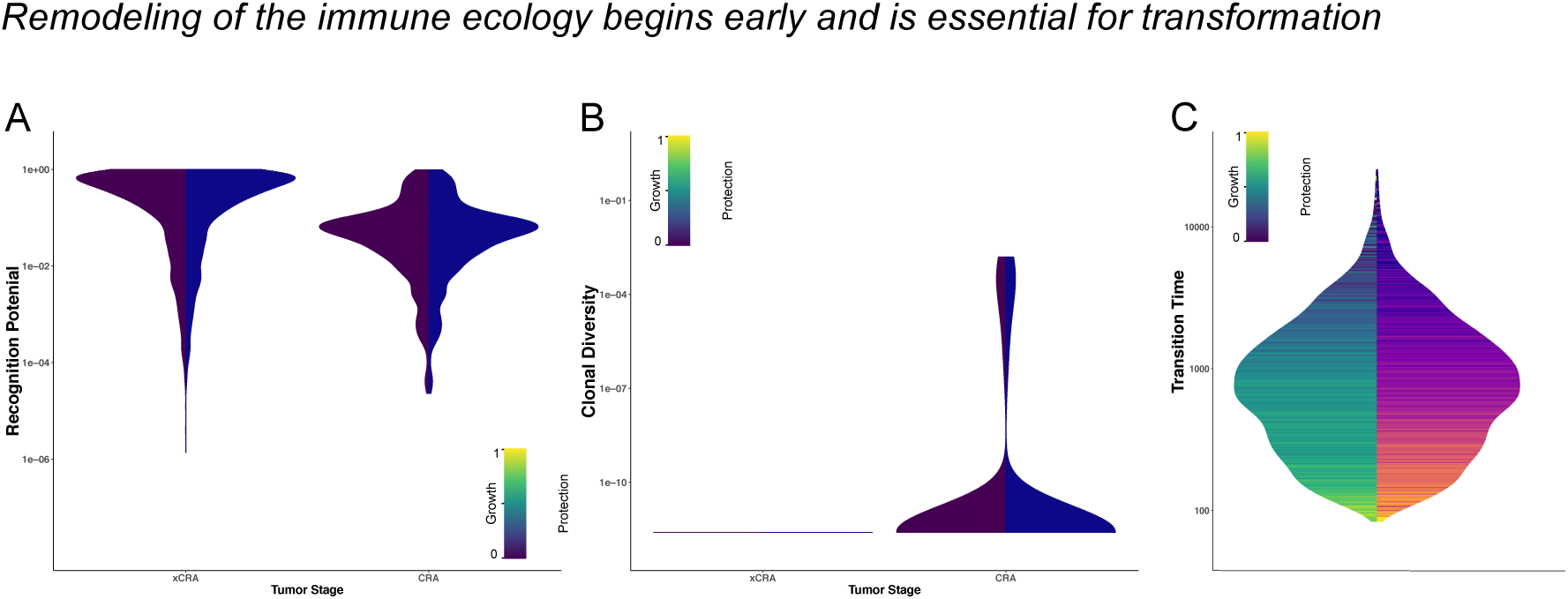
Immune ecology and aITH before immune escape. **A** Model-predicted distribution of neoantigen recognition potentials in the absence of immune escape, for adenomas that were eventually eliminated (xCRA) and those that successfully progressed to CRC (CRA). The color on the left side of the violin indicates the growth benefit received by the clone in which the antigen originated, i.e. the “source clone”, due to that clone’s recruitment of immunosuppressive cells. The color on the right side of the violin reflects how much the source clone was able to protect itself from attack by immune blockade and/or immune suppression. **B** Clonal diversity, calculated as 1-Simpson’s *D*, prior to immune escape. **C** The amount of time between adenoma formation and the foundation of the subsequent carcinoma, colored by the adenoma founder clone’s immune escape phenotype.

Higher efficacy of immune escape reduced the time required for evolution of malignancy (Figure 3C). Values of protection due to immune blockade, protection from suppression, and growth benefits from recruiting immunosuppressive cells varied across our simulations. Transition from CRA to CRC occurred most rapidly in tumors that were strongly protected from immune attack (due to the combined protection of blockade and immunosuppression) and that had high growth benefits associated with recruited immunosuppressive cells. Immune escape facilitates this rapid transition because new clones experience low immune predation, and the generation of more clones increases the probability that one will acquire the necessary driver mutations to become the founder of a carcinoma. This result suggests that adenomas exhibiting higher concentrations of immunosuppressive cytokines, growth factors, checkpoint inhibitors, etc. may be at a higher risk of rapidly progressing to carcinoma, due to lack of immune control.

The amount of ecological remodeling that occurs during immune escape determines patterns of aITH. Even when escape mechanisms are present, there remains some selection against antigens with high recognition potentials (Figure 4A). Any adenomas that do have such antigens and are only able to weakly mitigate attack, are rapidly eliminated by the predatory immune response (xCRA). In comparison, antigens with high recognition potentials can survive and progress to CRC if their source clone had the ability to protect itself from immune predation. CRA that have weak escape mechanisms must “get lucky”, as only clones that acquire antigens with low recognition potential survive immune surveillance.

**Figure 4.**
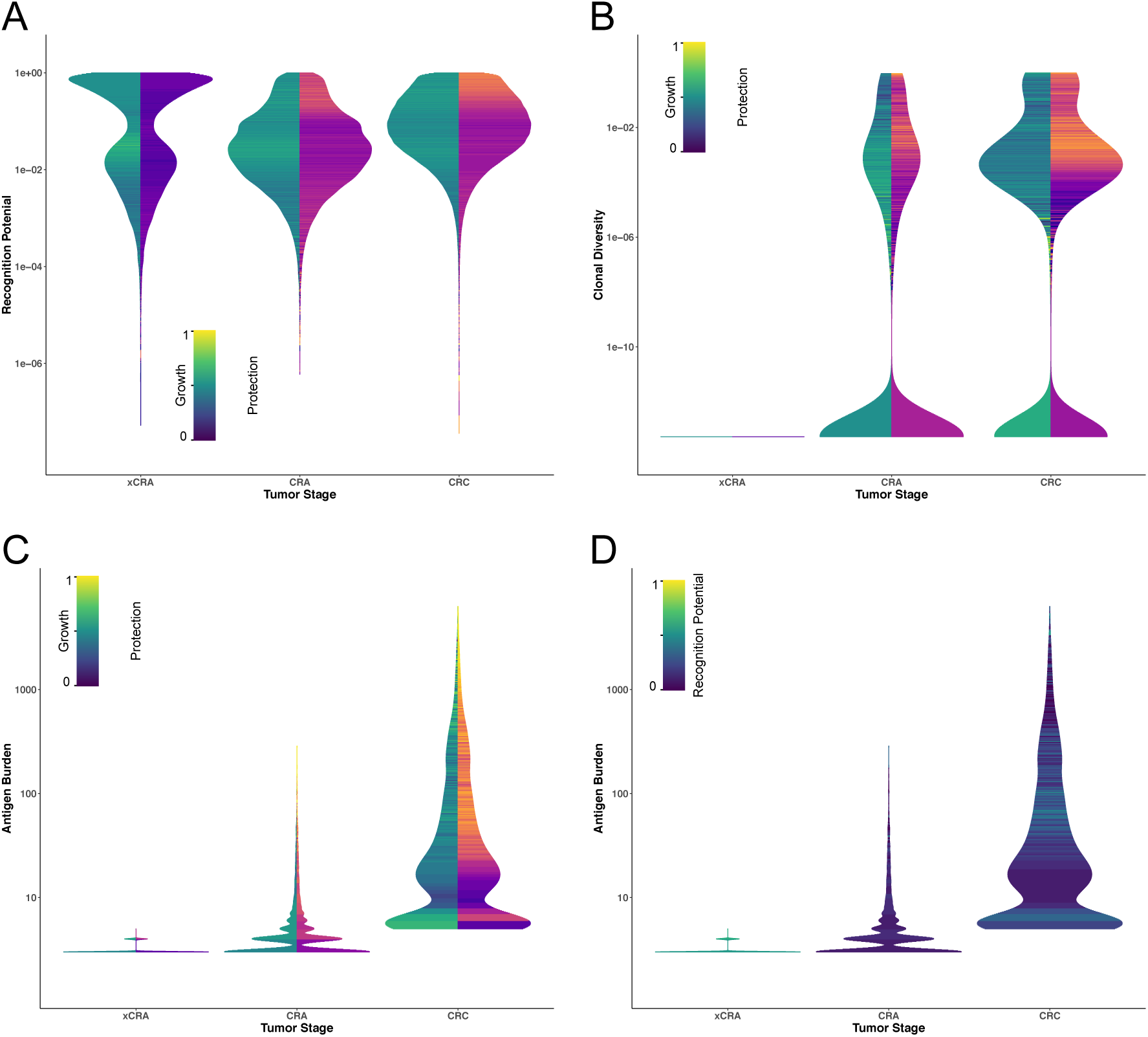
Tumor-immune ecology shapes aITH, with stronger immune escape leading to higher recognition potentials, higher antigen burden, and greater clonal diversity. **A** Model-predicted distributions of recognition potentials given a clone’s immune escape phenotype, for all xCRA, CRA, and CRC. The color of each horizontal bin reflects the average weighted phenotype of the antigen’s source clone that falls within that bin, where the weights are the source clone’s VAF. **B** Clonal diversity for all xCRA, CRA, and CRC. **C & D** Tumor antigen burden as a function of tumor stage, colored by the weighted average escape phenotypes (**C**) of clones within each tumor, or average weighted recognition potential (**D**), where the weights are the clone size in the tumor. ***Immune escape via ecological remodeling is a common phenomenon in colorectal cancer***

In Figure 4B, adenomas that are eliminated (xCRA) are characterized by low levels of diversity, as without protection, new clones that acquire a neoantigen and inherit ancestral antigens cannot survive immune predation, resulting in a monoclonal tumor. Conversely, as the strength of immune escape increases, so too does diversity, as there is weaker negative selection against new clones, which are always more antigenic than their ancestors. Figures 4C&D show the antigen burden, colored by phenotype (Figure 4C) and recognition potential (Figure 4D). Extinct adenomas are characterized by a weak immune escape phenotype and a handful of antigens that have high recognition potentials. On the other hand, adenomas that progress are characterized by a larger number of antigens, but selection against high recognition potentials remains strong. Carcinomas share these patterns, but in a more extreme form, i.e. compared to CRC that rely on passive escape, CRC with active escape have even higher clonal diversity and antigen burdens, being able to sustain more antigens with high recognition potentials.

### Immune escape via ecological remodeling is a common phenomenon in colorectal cancer

Simulation results suggest that immune escape, driven primarily by immune suppression, occurs early and is required for progression from adenoma to carcinoma. In the model, escape mechanisms are modeled as phenotypes, but those phenotypes alter the ecology by recruiting immunosuppressive cells and/or blocking immune attack. Thus, the model predicts that there exists succession in the tumor ecology during progression, with a transition from a “hot” T-cell rich ecology, to one that is “cold” and immunosuppressive. This pattern would be accompanied by an increase in antigen burden and the presence of antigens with higher recognition potentials, due to the lack of negative selection against highly antigenic mutations.

We examined primary human tumors for evidence of these patterns. Specifically, we considered colorectal adenomas (CRA), colorectal carcinomas (CRC), and “ca-in-ad” (CIA) samples, that latter of which consist of a nascent carcinoma (C-CIA) emerging from its precursor adenoma (A-CIA) (Figure 5). CIA samples not only allowed us to study an early stage of carcinogenesis, but they also provided a unique opportunity to directly compare a late adenoma and an early carcinoma within the same sample.

**Figure 5.**
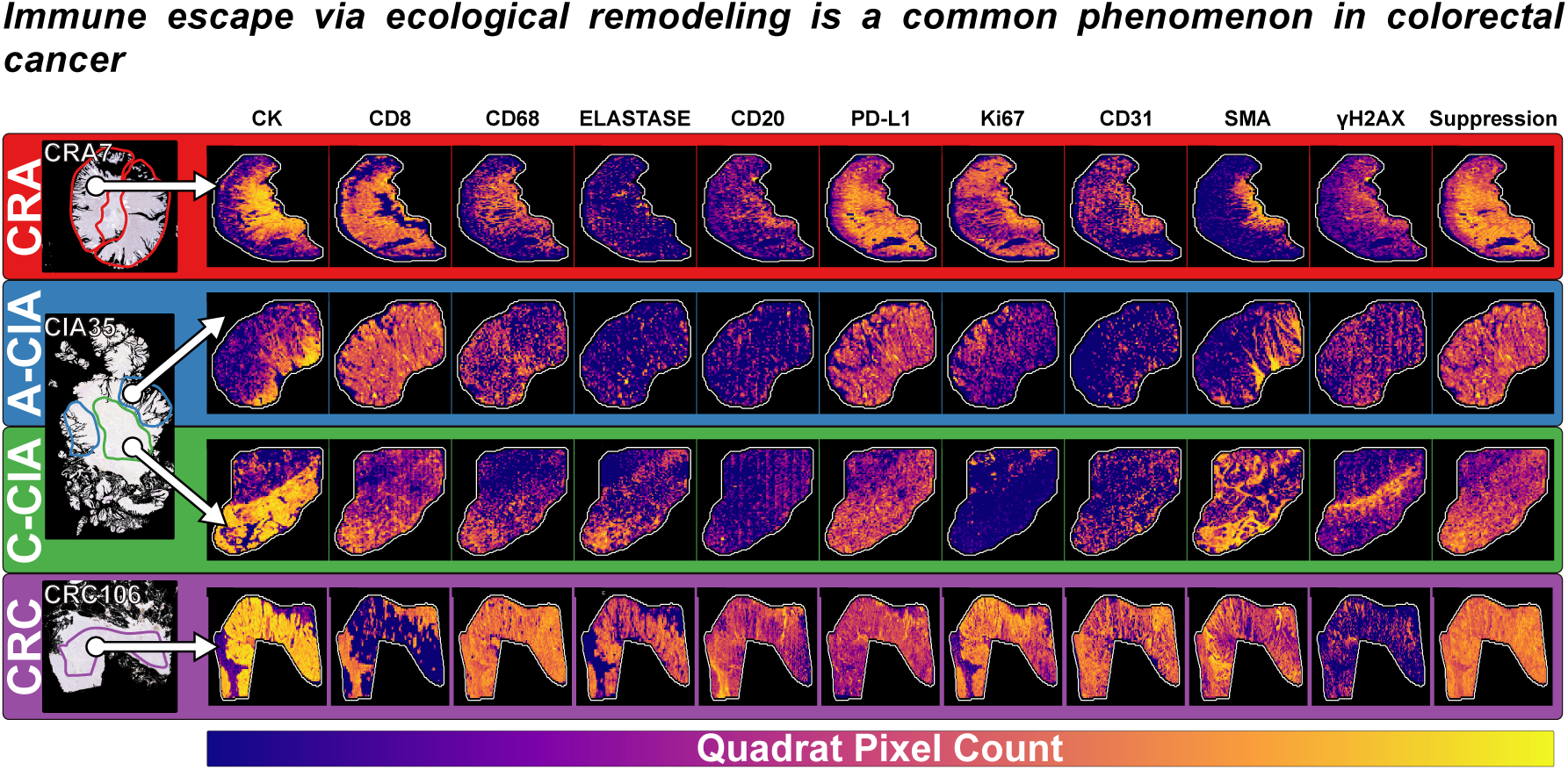
The tumor-immune eco-evolutionary status was characterized for a cohort of colorectal cancer tumors. There are three types of samples: adenomas (CRA), carcinomas (CRC), and “ca-in-ad” tumors (CIA), which are samples in which carcinomas (C-CIA) are emerging from a precursor adenoma (A-CIA). The tumor ecology was quantitatively described using 10 markers applied across a set of serial slices stained using multi-color immunohistochemistry. Stain segmentation of the set of aligned images created a composite image of the tumor ecology. Spatial interactions were determined by slicing the composite images into quadrats and then using the quadrat pixel counts to build a species interaction network. Here, the log of the quadrat pixel counts are shown for each of the 10 markers. The final “Suppression” column is the combined abundances of macrophages, neutrophils, and PD-L1. CK=epithelial cells, CD8=cytotoxic T cells, CD68=macrophages, Elastase=neutrophils, CD20=B cells, PD-L1=programmed death ligand 1, Ki67=proliferation, CD31=vasculature, SMA=activated fibroblasts, yH2AX=DNA damage.

We measured tumor ecology using whole slide multi-color immunohistochemistry (IHC) on tumor serial sections of CRAs, CRCs and CIAs (Figure 5). We performed whole slide image registration (WSIR) of serial multi-color IHC to quantify features of the tumor ecology (see Supplemental for examples of IHC). For this analysis, there were 12 CRA, 26 CIA, and 15 CRC.

Intra-tumour antigen heterogeneity (aITH) was measured using multi-region exome sequencing, with a downstream pipeline to determine neoantigen presentation and recognition potentials for 6 CRAs, 3 CIAs, and 7 CRCs. Sequencing was normalized for depth between samples. In total, 2 CRAs, 3 CIAs, and 3 CRCs had both imaging and genomic data. We tested for the temporal trends predicted by the model during the process of immune escape by assuming that the A-CIA adenoma will ultimately be out-competed by its emerging C-CIA carcinoma, resulting in the existence of a distinct CRC. In other words, we assumed the changes observed from A-CIA to C-CIA to CRC are similar to those that would occur during evolution from late adenoma to mature carcinoma. As CRA often remain benign (Hofstad et al., 1996; Logan et al., 2012; Zauber et al., 2012), they are excluded from temporal analyses, but are instead compared directly to A-CIA to determine how (currently) benign adenomas (CRA) differ from adenomas that have begun the transition to malignancy (A-CIA).

#### Describing tumor ecology using serial multi-color IHC

The tumor ecology can be described using IHC. We used multi-color IHC on thin serial sections to measure the abundance of various cell types (Tables S2-3): tumor (epithelial) cells (CK+), T cells (CD8+), macrophages (CD68+), neutrophils (elastase+), and B cells (CD20+). Immune blockade was quantified with PD-L1 staining. The microenvironment was described by labeling for activated fibroblasts (aSMA+) and vasculature (CD31+). In addition, IHC was used to examine proliferation status (Ki67 staining) and DNA damage (yH2AX staining). A mask was applied to each sample so as to isolate the region of interest, allowing us to measure cell abundance in only the adenoma and/or carcinoma region of the sample. While there are only two IHC markers per slice, we utilized WSIR to align the thin serial sections, creating a composite image that preserves the relative spatial distributions of each cell type. We then quantified spatial cell-cell interactions using a method based on partial correlations that takes indirect effects into account, something that can confound simple spatial correlation (Morueta-Holme et al., 2015). Stain segmentation was performed using a support vector machine (see Supplemental Methods for more details). All of the image analysis was performed on the full resolution images. Complex and spatially variegated immune cell composition was observed (Figure 5)

#### Ecological analysis of IHC reveals succession from a hot anti-tumor immune ecology to one that is cold and pro-tumor

Based on ecological analysis of the samples, benign adenomas (CRA) do not appear to have escaped immune control. Their ecologies are defined by the presence of CD8 T cells and PD-L1 as indicator species, i.e. cell types that uniquely reflect the state of the tissue, distinguishing that stage from all others (Figure 6A) (Caceres, 2013). This pattern is consistent with the simulated adenomas that go extinct (xCRA), in that these adenomas stimulate such a strong immune response that they are eliminated despite having some degree of moderate protection (Figure 4). CRC, on the other hand, are defined by tumor cells, vasculature, and potentially immunosuppressive neutrophils, each of which is an indicator species for that stage. Constrained analysis of principal coordinates (CAP) and permutational multivariate analysis of variance (PERMANOVA) (Marti J. Anderson, 2001; Marti J. Anderson & Willis, 2003; Oksanen et al., 2018) (Figure 6B) reveals a significantly different (p=0.001) makeup of tumor ecology of CRA, CIA, and CRC, suggesting succession through distinct ecologies as a the tumor progresses.

**Figure 6.**
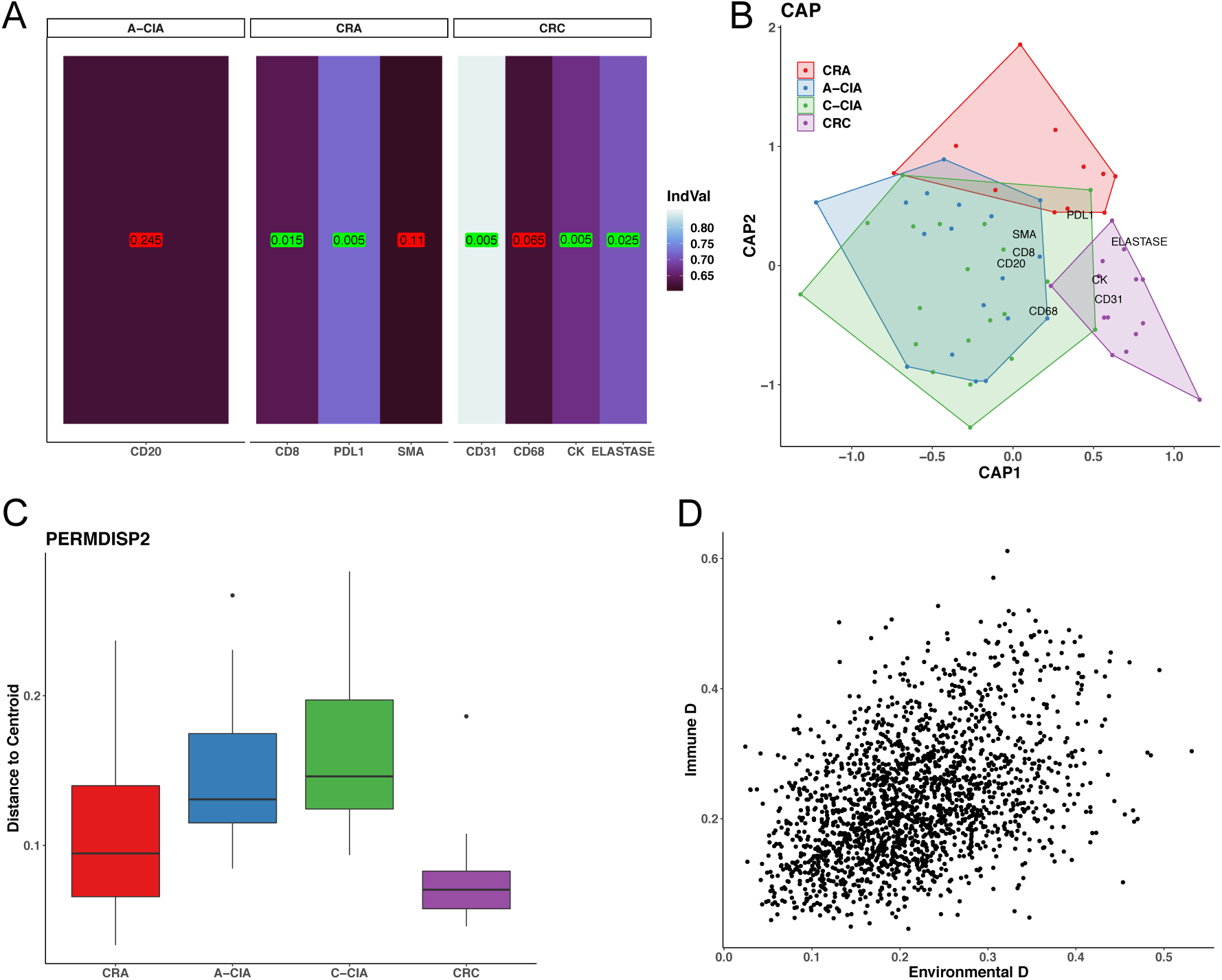
Tissue level ecological analysis reveals adenomas and carcinomas have unique and distinct cold and hot ecologies, respectively. **A** Indicator species analysis was performed to determine which, if any, cells define each tumor stage. The indicator value (IndVal), describes how strongly each cell defines the tumor stage. The ecologies of A-CIA and C-CIA do not have any significant indicator species. **B** PERMANOVA comparing ecologies across tumor stages, which can be visualized using constrained analysis of principal coordinates (CAP). **C** PERMDISP2 describes the amount of intra-stage ecological heterogeneity by comparing intra-group dispersions of ecological distances. **D** The Mantel test finds that there is a significant correlation between environmental distances (defined by vasculature, tumor, fibroblasts, and PD-L1) and the distances in immune cell communities (cytotoxic T cells, macrophages, neutrophils, and B cells) (p=0.001).

Interestingly, CRA and CRC show the least amount of ecological heterogeneity, as revealed by PERMDISP2 (M. J. Anderson, 2006) (Figure 6C), while the transitory CIA stage seems to have a variety of ecological markers. This increase in heterogeneity supports the hypothesis of transition from a tumor-centric approach that defends against a “hot’ immune response in CRA, into a niche-construction-based approach that “cools” the immune response in CRC which occurs at the point malignancy develops. The intermediary CIA stage would like rely on aspects of both approaches, until the mature CRC emerges and the tumor-centric protection is no longer necessary. A convergence on a common ecological remodeling strategy by CRC is likely the result of strong selection pressure imposed by the immune system.

In the model, changes in the immune response are initiated by the tumor, and therefore differences in the tumor microenvironment and immune composition should be correlated. We tested for this relationship using the Mantel test (Legendre & Legendre, 2012) to detect correlations between the ecological distances in immune-cell lineages (T cells, B cells, macrophages and neutrophils) and environmental factor distances (tumor cells, fibroblasts, PD-L1, and vasculature). Test results reveal that there is indeed a significant interrelationship between differences in immune composition and other aspects of the tumor ecology (p=0.001) (Figure 6D). Such a pattern is again consistent with hypothesis of eventual immune escape via immunosuppressive niche construction.

The model predicted that adenomas that aggressively progressed to carcinomas were more likely to have developed a component of immune evasion through niche construction (Figure 3). Using a combination of frequentist and permutation tests, we examined the trends in cell abundance between CRA and A-CIA samples (Figure 7A) and found that the adenomas from CIA samples (i.e. those that are known to have given rise to their adjacent carcinoma) have a decrease in cytotoxic T-cell abundance and an increase in macrophage presence, compared to the CRA samples that likely include many benign adenomas that would never progress. This is consistent with the hypothesis from the model that progressing adenomas will have a “colder” immune ecology at the time of transformation into carcinoma. Furthermore, PD-L1 decreases from CRA to A-CIA, suggesting less reliance on immune blockade and suggesting that PD-L1 expression is preventing CRA from being eliminated, but niche engineering through immune suppression is vital for adenomas to progress further.

**Figure 7.**
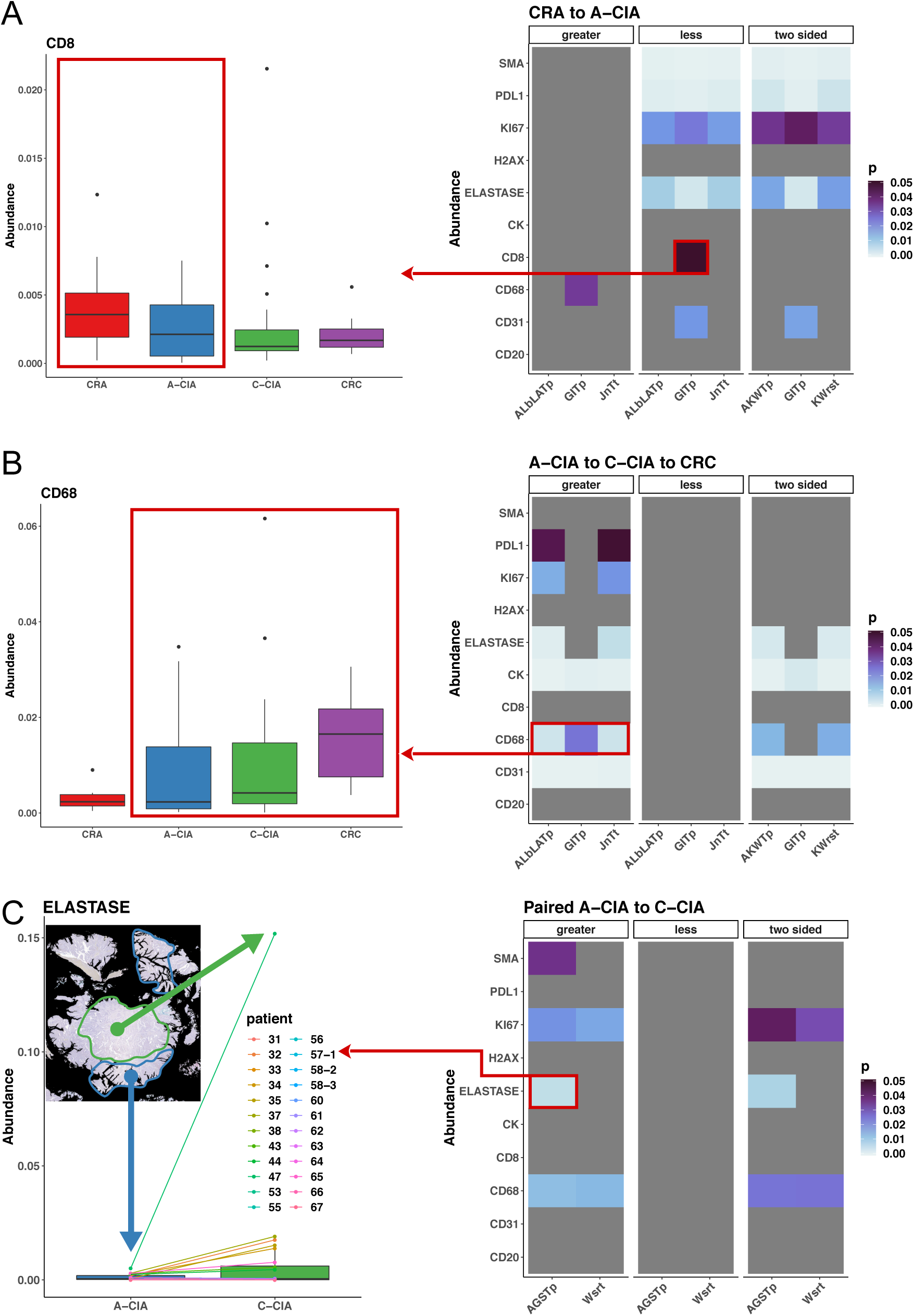
Changes in abundance consistent with ecological remodeling during immune escape. The abundance of each marker was determined using multi-stain IHC, where the abundance is considered to be proportional to the number of pixels that were positive for each stain. All values were normalized by dividing the pixel counts by the total area of the tissue, which was masked to include only the adenoma/carcinoma, and exclude transparent pixels (i.e. areas with no tissue). For each table, the three major columns represents an alternative hypothesis for a set of statistical tests (minor columns): “greater” indicates an increase in the relative abundance for the titular stages; “less” means there was a decrease in the abundance; “two sided” indicates there are significant differences, which may or may not be directional. The color of each tile represents the significance of the test, where grey values indicate non-significance. **A** Changes in abundance of each marker from CRA to A-CIA. As an example, the left boxplot shows how the abundance of CD8 T cells significantly decreases from CRA to A-CIA, as indicated by the GITp test. **B** Assuming that carcinomas will always outcompete their precursor adenoma, changes in abundance during progression were estimated by testing for trends going from precursor adenoma A-CIA to the descendant C-CIA carcinoma and into a mature CRC. The boxplot on the left shows how the abundance of macrophages (CD68) increases with progression. **C** Intra-tumor changes in abundance were determined by using paired tests to compare each C-CIA to its own precursor A-CIA adenoma. The example paired plot on the left shows how the abundance of neutrophils (Elastase) is higher in each C-CIA compared to its ancestral A-CIA. KWrst= Kruskal-Wallis rank sum test, AKWTp=Approximative Kruskal-Wallis Test (permutation), GITp= General Independence Test (permutation), ALbLATp= Approximative Linear-by-Linear Association Test (permutation), JnTt= Jonckheere-Terpstra test.

To investigate the trajectory of adenomas to full-fledged carcinomas, we assume that the progression from A-CIA to C-CIA to CRC is representative of what occurs in patients when a late adenoma becomes a full mature carcinoma. Consistent with the model’s prediction that the process of escape begins early and continues through progression, we see significant increases in potentially immunosuppressive macrophages (CD68) and neutrophils (elastase), as well as PD-L1, consistent with what one would expect during a continued shift to an immunosuppressive ecology (Figure 7B).

We also examined the pairwise ecologies of each A-CIA and its emerging C-CIA within each CIA sample (Figure 7C) using paired statistical tests. As in the cohort analysis, we see a consistent trend across all CIA samples that the carcinoma, compared to its neighboring precursor adenoma, has significantly higher abundances of macrophages, neutrophils, and proliferating cells (Ki67). These intra-tumor trends provide further evidence that carcinomas are remodeling their ecology to create an immunosuppressive environment.

Given the spatial information present in the samples, we created species interaction networks (Blonder & Morueta-Holme, 2017; Morueta-Holme et al., 2015), using the composite images generated by WSIR. The weights of each pairwise cell-cell interaction were then used to test if the spatial distribution of cells further supports the hypothesis of ecological succession. Figure 8A shows a representative CRC sample stained for tumor cells (CK), cytotoxic T cells (CD8), and neutrophils (elastase), showing that the neutrophils have infiltrated the tumor, while the T cells are spatially segregated. Indeed, for the set of CRC samples (Figure 8B), cytotoxic T cells and tumor cells are found in different areas, while tumor cells and PD-L1 are frequently found together. As in Figure 7, we examined the changes in spatial interactions arising during progression from A-CIA to C-CIA to CRC (Figure 8C). Among the many changes in interactions, the most relevant to immune remodeling are the increases in interactions between tumor cells and PD-L1; and tumor cells and neutrophils. Decreases were observed in the interactions between cytotoxic T cells and tumor cells; and between neutrophils and cytotoxic T cells. This supports the hypothesis of a shift from hot to cold tumor ecologies, given an apparent breakdown of T-cell infiltration and engagement of the tumor cells. It is worth noting that while the overall abundance of cytotoxic T cells did not change significantly during progression, their spatial localization with respect to tumor cells did change significantly. This highlights the importance of taking spatial distributions into account when investigating the ecologies of each stage of progression.

**Figure 8.**
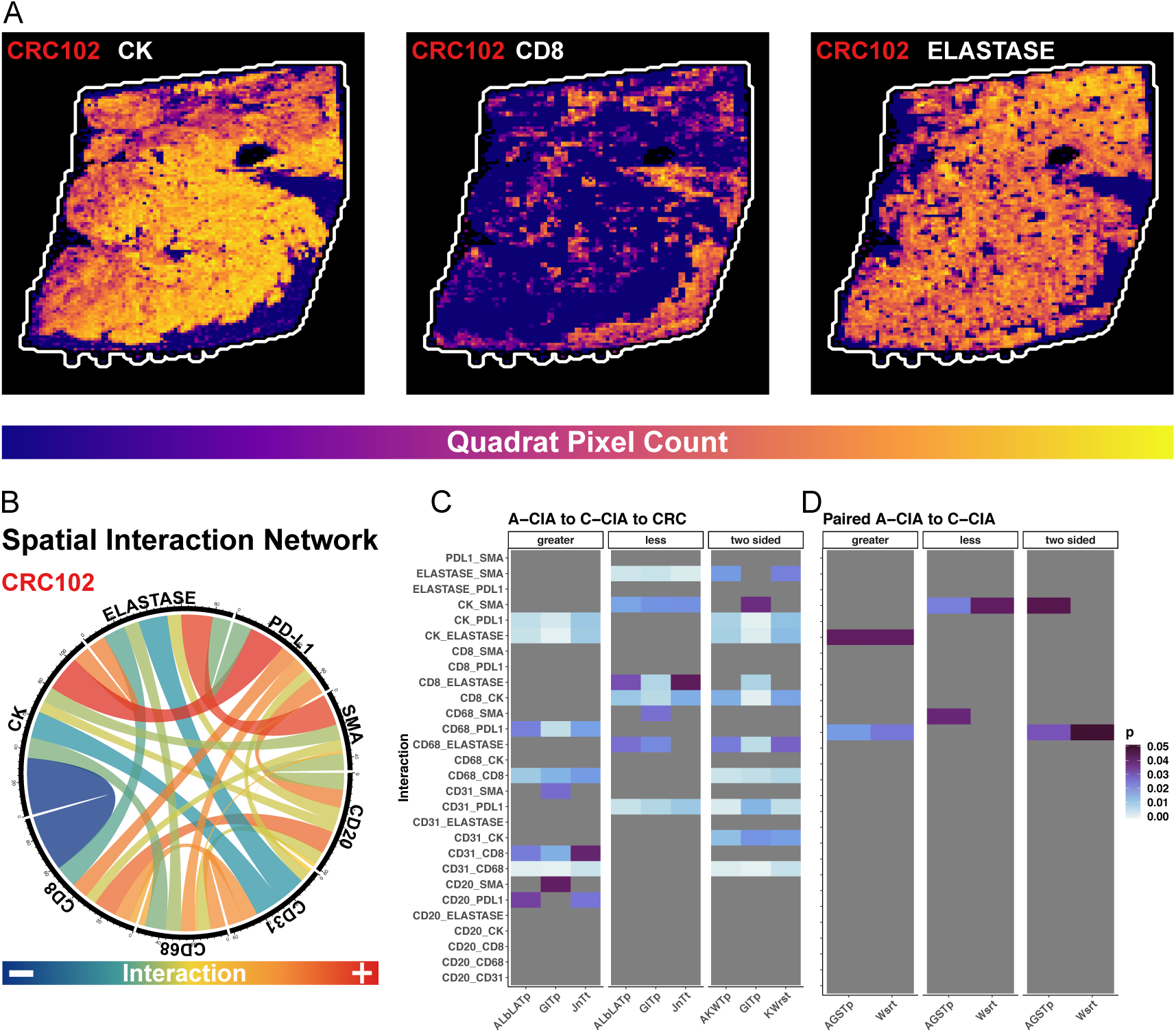
Spatial analysis of interactions reveals a temporal succession from a hot immune ecology to one that is cold. For each sample, the series of multi-stained IHC images were aligned and divided into quadrats. The abundance of marker in each quadrat was then determined. This process creates a composite image of the tumor ecology, with each quadrat containing the abundances of multiple cell types. The quadrat counts can then be used to visualize the spatial distributions and to construct interaction networks. **A** Spatial segregation between cell types within a representative CRC sample. **B** Spatial species interaction networks were constructed using the quadrat counts. In this example chord diagram, the links indicate interactions, the width of each link is proportional to the strength of the interaction, and the color defines the direction of the interaction, where negative means the two cell types are found in different areas, and positive values indicate the two cell types are found together, after having taken indirect effects into account. **C** Changes in interactions assuming changes during progression are similar to the changes from A-CIA to C-CIA to CRC. Greater (less) indicates an increase (decrease) in the interaction during progression. **D** Paired tests were used to detect intra-tumor changes in cell-cell interactions, by comparing A-CIA adenomas to their descendant C-CIA carcinomas.

While not as varied, some evidence of niche construction is evident in pairwise analysis of A-CIA and C-CIA within each CIA sample. As with the changes observed during progression of CRA to CRC, the interaction between tumor cells and neutrophils is significantly stronger in C-CIA than within their precursor A-CIA. This, again, suggests that immune remodeling is occurring within the nascent carcinoma.

Taken together, these data indicate the generation of significantly different ecologies across tumor stages, and are consistent with the changes the model suggests we should observe before, during, and after active immune escape. Benign CRA stimulate a strong T-cell response and lack immune suppression, and thus appear to remain under immune control, if not removed altogether. However, recruitment of, and interactions with, immunosuppressive cells continue to increase through progression from late adenoma (A-CIA) to mature carcinoma (CRC), while there is a simultaneous drop in immune predation, generating an ecology supportive of tumor growth.

### Observed patterns of antigenic intra-tumor heterogeneity consistent with immune escape

The eco-evolutionary dynamics of immune predation and escape shape aITH. Simulations reveal that antigen burden and the distribution of recognition potentials are good indicators of whether or not the immune system is successfully responding to the tumor (Figure 9A). In the absence of immune suppression or blockade, antigen burdens are small and recognition potentials are low. Conversely, if active immune evasion has occurred, selection against immune recognition is relaxed, and both burdens and the recognition potentials are free to increase.

**Figure 9.**
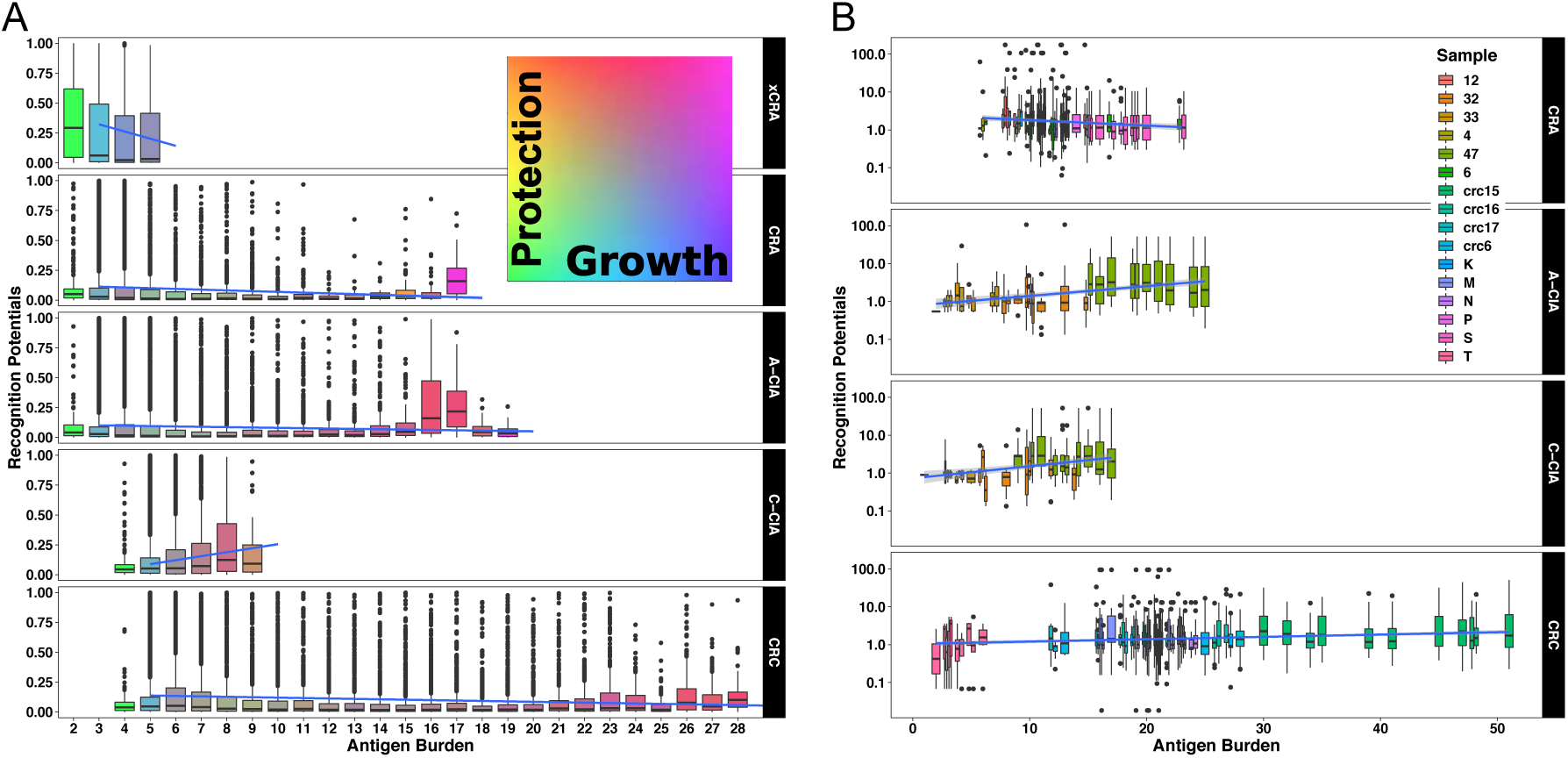
Observed patterns of aITH are consistent with active immune escape. **A** The distribution of recognition potentials within simulated tumors with each antigen burden level. The color of each boxplot indicates the average escape phenotype of the tumors with that particular burden, where the tumor phenotype is defined as the average clone phenotype. Blue lines are the trend lines for the relationship between burden and recognition potential. **B** The observed distribution of recognition potentials versus antigen burden in samples, with trend lines shown in blue. Due to unequal sequencing depths, each sample underwent multiple replicates of downsampling. The distribution of recognition potentials within each burden per replicate are shown.

Among the adenomas from patient samples, there are several samples that have low burdens and high recognition potentials, versus one that has a higher burden and low recognition potentials (Figure 9B). As these lesions were resected, it is impossible to know which ones would have progressed. However, the pattern of aITH of the former is similar to those simulated adenomas that were eliminated (xCRA, Figure 9A), having low burdens but high recognition potentials, thus suggesting that these samples may have been subject to strong predation and likely to be controlled, if not eliminated, by the immune system. However, CRA sample S has a higher burden and lower recognition potentials, which more closely resembles adenomas that had progressed (CIAs). It is noteworthy that within both observed adenomas and those that go extinct in simulations, there is a negative relationship between burden and recognition potential, which is again consistent with simulated tumors that have not yet mitigated immune attack, as there remains strong negative selection against acquiring antigens with high recognition potential. It is possible that adenoma S was persisting due to passive escape.

Overall, the observed patterns of antigen burden and the distribution of recognition potentials from A-CIA to C-CIA to CRC are consistent with what one expects under increasing development of active immune escape (Figure 9). In particular, the antigen burden of CRC is significantly higher than the other tumor stages (Tukey’s HSD for CRC vs. CRA p=7.7e-9, CRC vs. A-CIA p=2.3e-7, CRC vs. C-CIA p=3.3e-12). Compared to CRA, CRC also have significantly more antigens with higher recognition potentials (Kolmogorov-Smirnov test p=0.01). In Figure 9B, it is striking that the observed C-CIA samples have lower antigen burdens than their precursor A-CIA adenomas. However, this same pattern is observed in the simulated tumors (Figure 9A), and is due to the carcinomas emerging from an adenoma clone. As these nascent carcinomas are recent descendants of an adenoma, insufficient time had elapsed to generate clonal diversity within the carcinoma component.

In simulated tumors, directional selection against antigens with high recognition potentials relaxes as the strength of immune escape increases, facilitating the existence of tumors with high antigen burdens. This is visible in Figure 9A, which shows that the strong negative relationship between recognition potentials and antigen burden observed in adenomas that go extinct (xCRA) weakens during progression, evidenced by the trendline that switches from being negative to one that is more flat. However, in simulations, lower recognition potentials are *always* advantageous to tumour cells, as immune attack is never completely alleviated. It is therefore remarkable that in patient samples, the observed negative relationship between burden and high recognition potential in CRA becomes positive in tumors that have transformed (CRC), or are in the process of doing so (CIA) (Figure 9B). This pattern suggests that selection against high recognition potentials is dramatically reduced during progression, in other words that cancers have evolved the ability to (near completely) escape immune predation.

## Discussion

Our analyses reveal that immune evasion is fundamental in the evolution of benign colorectal adenomas to malignant carcinomas. CRAs are characterized by high cytotoxic T cell abundance, low macrophage abundance, low antigen burden, and high recognition potentials. This ‘hot’ ecology and pattern of aITH is consistent with simulated adenomas that lacked escape mechanisms and were eliminated by immune predation. We thus conclude that these (currently) benign CRA have yet to break out of the immunogenic bottleneck.

Adenomas that are known to have progressed (i.e. A-CIA) have initiated the process of remodeling the ecology to suppress immune attack, a process that continues from nascent to mature CRC. This is evidenced by increasing abundances of macrophages and neutrophils, tumor interactions with neutrophils and PD-L1, all of which is accompanied by decreases in cytotoxic T-cell interactions with tumor cells. The result of this process is the generation of a ‘cold’ ecology, defined by tumor cells, neutrophils, and vasculature, an ecology shared by all CRCs. Modeling predicts that under such high immune suppression, antigen burdens should be large and high recognition potentials are tolerated, and indeed, the patterns of aITH observed in clinical samples fit this pattern. Taken together, these results strongly suggest that passage through the immunogenic bottleneck that facilitates evolution of CRC begins early, and is driven by immunosuppressive niche engineering.

Neutrophils appear to play a particularly important role in engineering the cold CRC ecology. Their abundance increases not only through CRC progression, but also from the adenoma to the adjacent carcinoma region within the same tumor (i.e. A-CIA to C-CIA). The same can be said for their interactions with tumor cells. In addition to vasculature and tumor cells, presence of neutrophils also distinguishes CRC from CRA.

Analysis of IHC highlights the importance of considering spatial context when examining tumor-immune interactions. Despite the fact that there are not significant differences in the abundances of cytotoxic T cells during progression, there is clear spatial segregation of cytotoxic T cells and tumor cells in later stages. Only by analyzing the spatial structure does it become clear that cell-cell interactions between tumor cells and cytotoxic T cells decrease during progression, allowing later-stage tumors to effectively avoid predation.

The conclusion that niche engineering is initiated early and facilitates passage through the immunogenic bottleneck has important clinical implications. Stimulating immune predation may be ineffective in the presence of immunosuppressive cells and/or if there is spatial segregation of tumor cells and cytotoxic T cells. Indeed this may explain why checkpoint inhibitors and DC vaccines, which are intended to enhance cytotoxic T-cell attack, have limited efficacy in microsatellite stable (MSS) CRC (Kalyan, Kircher, Shah, Mulcahy, & Benson, 2018; Passardi, Canale, Valgiusti, & Ulivi, 2017). Instead of aiming to re-activate T cells, it may be more effective to re-engineer the ecology to be hot, possibly by re-polarizing immunosuppressive cells, a treatment currently being explored for macrophages (Brown, Recht, & Strober, 2017; Genard, Lucas, & Michiels, 2017; Kulkarni et al., 2018).

A second clinical implication of niche engineering being initiated early is that the tumor ecology and pattern of aITH present in CRA may be a prognostic marker for progression. Modeling suggests that the rate of progression from CRA to CRC is determined by the tumor ecology. Simulated CRA with hot ecologies progressed more rapidly than those that had colder ecologies. One might thus expect that a CRA with a hot ecology, large antigen burden, and high recognition potentials, would be at a higher risk of progressing than a CRA that is cold and has a low antigen burden. If immune ecologies are correlated between multiple adenomas that arise in patients, then measurement of intra-adenoma immune ecology could determine a patient’s future risk of developing CRC.

The eco-evolutionary model provides insights into the status of tumor ecology and aITH under varying degrees of immune escape. While most CRC appear to have undergone extensive immunoediting during the transition from CRA, the model also describes tumor ecology and aITH when escape mechanisms are weak or non-existent. We propose that a patient’s tumor ecology and aITH can be compared to these predicted patterns to get a sense of how protected the tumor is from immune predation. For example, a CRC sample with a pattern of aITH that is consistent with passive escape (i.e. avoiding elimination by having antigens with low recognition potentials) may have a potentially effective anti-tumor immune response, and may therefore be a good candidate for immunotherapy that increases the efficacy of cytotoxic T cell attack. In this way, the simulation results may be of clinical utility in guiding treatment decisions through measurement of tumor ecology and aITH.

In summary, we provide evidence of a critical role for immune predation in preventing colorectal malignancy, and further the necessity of evolving immune escape mechanisms is the key bottleneck in disease progression.

## Author Contributions

CDG developed and analyzed the computational model, developed the methods to segment and align the immunohistochemistry, and performed analysis of the data collected from immunohistochemistry and neoantigen predictions. AMB optimized and performed all laboratory experiments. ROS processed whole exome sequencing of samples and downstream data generation and WCHC, PM and AS provided additional support in genomic analysis. MPN and SYH assisted AMB in laboratory analyses. MRJ, SL and AS graciously provided samples and/or data. MRJ and MJ provided histopathological assessment. MRT, TAG, and ARAA guided the research and supervised the writing of the manuscript. TAG and ARAA conceived the study.

## Acknowledgements

CG, MRT and ARAA are supported by Physical Sciences Oncology Network (PSON) grant from the National Cancer Institute (grant no. U54CA193489). MRT and ARAA are also supported the Cancer Systems Biology Consortium grant from the National Cancer Institute (grant no. U01CA23238). ARAA would also like to acknowledge support from the Moffitt Cancer Center of Excellence for Evolutionary Therapy. TG is supported by the Wellcome Trust (202778/Z/16/Z) and Cancer Research UK (A19771). AS is supported by the Wellcome Trust (202778/B/16/Z) and Cancer Research UK (A22909). We acknowledge funding from the National Institute of Health (NCI U54 CA217376) to AS and TAG. This work was also supported a Wellcome Trust award to the Centre for Evolution and Cancer at the Institute of Cancer Research (105104/Z/14/Z). TAG and SL are grateful for support from the Bowel and Cancer Research Charity (project grant scheme). TAG received support from the Barts Charity (pilot grant scheme). SL was supported by Wellcome Trust Senior Clinical Research Fellowship (206314/Z/17/Z).

## Supplemental Methods

### Sample collection and processing

FFPE samples (n = 54) representing adenomas (CRA, n = 13), adenomas with foci of cancer (‘ca-in-ads’, CIA, n = 24) and carcinomas (CRC, n = 17) were selected from the histopathology archives of University College Hospital, London, under UK ethical approval (07/Q1604/17) or John Radcliffe Hospital, Oxford under ethical approval (10/H0604/72). From each block, 7 serial sections were taken at 4 micron thickness, the first was stained with haematoxylin and eosin (H&E) and used for histopathological classification by two expert pathologists (MRJ and MJ). The remaining 6 sections were used for dual color immunohistochemical staining. A further 6 sections at 5 micron were taken from a subset of blocks (n = 10), and used for DNA extraction.

### DNA extraction

Different histopathological regions within individual lesions were demarcated on H&E slides by an experienced GI pathologist. This was used to guide careful needle dissection and DNA was then extracted from these discrete areas using DNA QIAamp Mini Kit (Qiagen) and standard protocols.

### Whole exome sequencing

Multi-region whole exome sequencing (WES) was performed on a subset of samples representing CRAs (n = 4 with two regions each), CIAs (n = 3, with one region from the carcinoma region and two from the adenoma region) and CRCs (n = 3, with two regions each). The quality of extracted FFPE DNA was verified using a multiplex PCR as previously described (van Beers et al., 2006) and only DNA samples that showed successful amplification of fragments greater than 300 bp in length were considered for WES. DNA input of 50ng was used to prepare sequencing libraries with the Nextera Rapid Capture Exome kit, according to manufacturers instructions (Illumina, Cambridge, UK). Libraries were sequenced with a target depth of 60x on Illumina’s HiSeq 2500 with 125 bp paired end reads (v4 chemistry). Additional samples were added from (Sottoriva et al., 2015) adding to the total number of multi-region CRAs and CRCs (n = 3 and n = 3, respectively). Alignments to the hg19 reference genome were conducted using the BWA-mem algorithm (Li & Durbin, 2010) and processed using the GATK best practices workflow (Van der Auwera et al., 2013) for downstream analysis.

### Variant Calling

Prior to calling variants, we normalized binary alignment/map (BAM) by down sampling to the lowest observed average depth across samples. First we calculated the average depth for target capture regions using samtools (Li et al., 2009). Once the average depth was calculated for each sample, the proportion of reads needed for each sample to reach the minimum average depth was calculated. Down-sampling was then performed using this proportion for each BAM file using PICARD (Broad Institute, 2017). This was conducted on each sample’s region to generate ten replicate BAM files. Variant calling was then performed for each replicate sample group (multiple regions against normal) using multiSNV (Josephidou, Lynch, & Tavaré, 2015), a joint calling method specifically designed for multi-region same patient experimental designs. Criteria used for assessing variant candidates dictated a minimum mapping quality of 30, a minimum base quality of 20, with at least five reads in the tumor and normal regions, and two variant alleles. Once a variant call set was obtained variants were further scrutinized for additional criteria. A total of ten reads were required within all normal sites and variant sites with no variant alleles present in the normal. Furthermore, a variant must be supported by a minimum number of two variant reads for at least one region; while the minimum variant allele frequency in one region is 0.1.

### Neoantigen Predictions

Human leukocyte antigen (HLA) haplotypes (A, B, and C) were called using PolySolver (Shukla et al., 2015) prior to down sampling on all normal regions for each patient. Neoantigen predictions were performed using NeoPredPipe (Schenck, Lakatos, Gatenbee, Graham, & Anderson, 2018), which utilizes ANNOVAR (Wang, Li, & Hakonarson, 2010) for variant annotations and netMHCpan (Jurtz et al., 2017), NeoPredPipe is specifically designed to handle multi-region sequence samples. Only MHC-class I neoantigens were assessed for peptides of 8, 9, and 10-kmer lengths. A minimum cut-off of 500nM binding affinities were used to be considered a putative neoantigen. To assess T-cell receptor binding potential an implementation of Łuksza *et al*.’s (Łuksza et al., 2017) recognition potential algorithm implemented within NeoPredPipe was used.

### Dual colour immunohistochemistry (IHC)

Sequential dual colour IHC of 10 markers was performed according to standard protocol. Briefly, 4µm serial sections were dewaxed, rehydrated and immersed in 3% hydrogen peroxide for 20 minutes to quench endogenous peroxidase activity. Antigen retrieval was carried out at 95°C for 20 minutes in sodium citrate buffer (pH 6.0), unless otherwise specified (Supplementary Table 2). After cooling, sections were incubated with blocking buffer (PBS supplemented with 2% goat serum and 1% bovine serum albumin) for 1 hour at RT. Primary antibodies were diluted in blocking buffer and applied for 1 hour at RT or overnight at 4oC (see Supplementary Table 2 for antibody details). Sections were then incubated with a biotinylated secondary antibody at RT for 45 min, followed by incubation with streptavidin-biotin peroxidase solution at RT for 45 min. Visualization of the first antibody binding was carried out using DAB, according to the manufacturer’s instructions (Vector Labs, Peterborough, UK). Slides then underwent a second round of antigen retrieval, generally at 95°C for 5 minutes in sodium citrate buffer (pH 6.0), before applying the blocking buffer for a further 1 hours at RT. The second primary antibody was then applied (see Supplementary Table 2 for details), followed by incubation with a biotinylated secondary antibody at RT for 45 min and incubation with streptavidin-alkaline phosphatase. Visualization of the second antibody binding was performed using Fast Red, according to the manufacturer’s instructions (Abcam, Cambridge, UK). Finally, sections were lightly counterstained using Gill’s haematoxylin, and allowed to dry before mounting and digitizing using the Pannoramic 250 high throughput scanner (3D Histech, Budapest, Hungary).

### Computational image analysis

All image processing was conducted using OpenCV (Bradski, 2000), scikit-image (Walt et al., 2014), and scikit-learn (Pedregosa et al., 2011; Schreiber et al., 2011) for the Python programming language.

#### Stain segmentation using support vector machine

Reliably separating brown (DAB) from red (Fast Red) is a challenging task, and so a support vector machine (SVM) was trained to estimate the probability that a pixel is positive for DAB, Fast Red, or is background. Four features were used to train and predict: the pixel’s RGB values, and the stain color descriptor (SCD) (Khan, Rajpoot, Treanor, & Magee, 2014). Three individuals performed the staining, and a SVM was created for each individual. In all cases, the accuracy of each SVM is above 90%.

Each image was sliced into 1000×1000 windows, and for each pixel in the window the SVM estimated the probability that the pixel is positive for each of the three stains. The SVM’s prediction was only accepted if the probability of correct classification was above 0.95.

Pixel classification generated a mask that indicated whether or not the pixel is positive for that stain. Once the stains in each window were segmented, they were stitched back together to create the full size image of the segmented stain. For each dual-stained image there were thus two full sized masks. The process was repeated for each image, generating 11 masks indicating positivity for each marker.

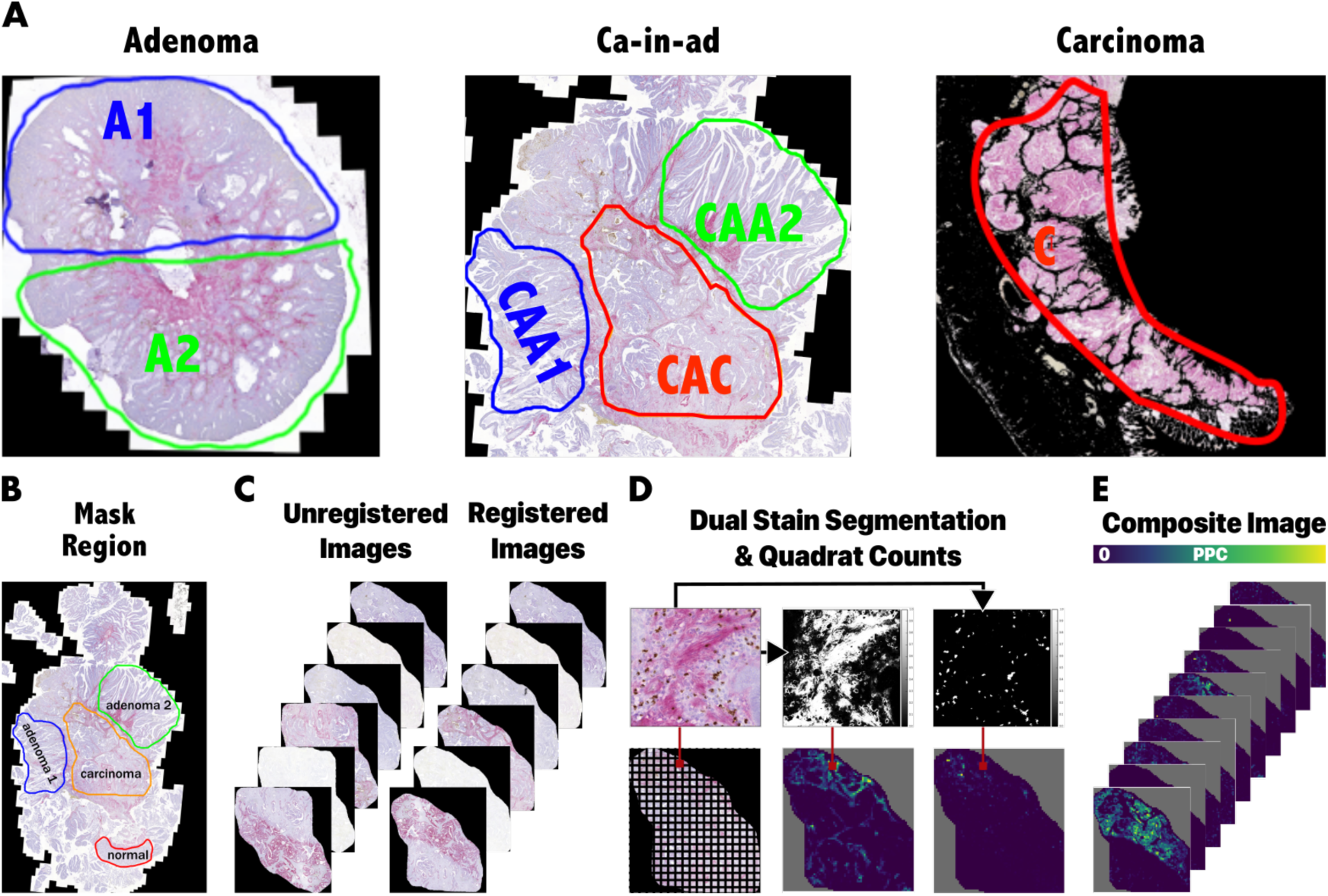
A) A cohort of colorectal cancer tumors were examined for the presence of immune escape. There are three types of samples: adenomas, carcinomas, and “ca-in-ad” tumors, which are samples in which carcinomas are emerging from the (presumably) precursor adenoma. Analysis of the tumor ecology was conducted using whole slide image registration performed on a set of dual stained serial sections for a total of 10 markers. Neoantigen recognition potential was predicted using from multi-region whole exome sequencing. B) Each sample has multiple regions that are analyzed independently by masking the area in the original image. C) Whole slide image registration is conducted on the entire set of IHC images, such that after registration the images are aligned with one another. D) A support vector machine was used to predict the probability each pixel is positive for either DAB or Fast Red. Classification was only accepted if the probability was greater or equal to 0.95. E) The transformation matrices found during registration were used to warp the segmented images so that they aligned, creating a composite image of the tumor ecology. The composite image was then sliced into quadrats and analyzed using a variety of tests from ecology.

#### Whole Slide Image Registration

The goal of image registration is to align all of the sample’s images to one another, generating a composite image in which all markers are correctly positioned relative to one another. To accomplish this, we determined, for each sample, which marker in the set of images had the most features in common with the other images in the set. The image with the most common features was used as the training image, and only the features in the query images that were common to all images were used to find the transformation matrices. The features detected in each image were KAZE features (Alcantarilla, Bartoli, & Davison, 2012), and matching was accomplished using brute force.

For each image, the matched features were used to find the affine transformation matrix, *A* _*i*_ that would to align each query image with the training image. Following the rigid affine transformation, enhanced correlation coefficient maximization (ECC) was used to find the homography matrix, *H* _*i*_, which would better align each query image to the training image (Evangelidis & Psarakis, 2008).

*A* _*i*_ and *H* _*i*_ were found using processed images that had been resized to have dimensions less than 1000 pixels in width and/or height. Therefore, the transformation matrices were scaled such that they could be applied to the full size images, which often have dimensions greater than 20,000 pixels in width and/or height. The transformations were applied to the images of segmented stains. This process yielded a composite image, in which each channel is a mask defining which pixels are positive for that marker. This composite image was then used to quantify direct cell-cell interactions.

#### Ecological Image Analysis

Description of the tumor ecology within and across various stages was accomplished using a variety of methods. Cell abundances (assumed to be proportional to the number of pixels positive for each stain) were measured from the un-registered images, normalized by tissue size, and then used to determine if there were directional changes over time, i.e. if there were significant increases or decreases in abundance during progression from late adenoma to mature carcinoma. The significance of these trends was determined using a combination of frequentist and permutation statistical tests. Using the registered composite images, a similar analysis was conducted to determine if there are directional changes in spatial interactions between the various cell types. This was accomplished by first dividing the composite image into quadrats, and then counting the number of positive pixels in each quadrat. The quadrat counts were then used to construct a species interaction network (Blonder & Morueta-Holme, 2017; Morueta-Holme et al., 2015), whose coefficients were compared across stages. In addition to testing for these interstage trends, paired tests were used to determine if there were significant intra-tumor changes in cell abundance and interactions. This was accomplished by comparing the carcinoma (C-CIA) to its precursor adenoma (A-CIA) in the same CIA sample.

Several ecological tests were used to compare tumor-immune ecologies across stages. We quantified and compared the amount of ecological homogeneity within each tumor stage, using multivariate homogeneity of group dispersions (PERMDISP2)(M. J. Anderson, 2006). Permutational multivariate analysis of variance (PERMANOVA) was used to determine if there were significant differences in the structure of the ecological communities of each tumor stage, which was then visualized using constrained analysis of principal coordinates (CAP) (Marti J. Anderson & Willis, 2003; Oksanen et al., 2018). Using indicator species analysis, we determined which, if any, cell types define the different tumor stages (De Cáceres & Legendre, 2009). Finally, the Mantel test was used to determine if differences in the immune ecology and microenvironment are correlated (Legendre & Legendre, 2012), as might be expected to occur during tumor instigated immune remodeling.

The types of analyses of each composite image can be divided into two categories: tissue-wide, in which the positive pixel counts (PPC) for the whole region was used; and spatial, in which the PPC per quadrat were used. For the tissue wide analysis, there were 12 adenoma, 24 ca-in-ad, and 15 carcinoma samples for markers H2AX, CD68, CK, Elastase, CD20, Ki67, SMA, CD8, and CD31. For PD-L1 there were 11, 18, and 14 samples respectively. Not all image alignments were successful, so there were fewer samples used in the spatial analysis. For the spatial interactions, only samples that had all markers aligned were used, and there were 9, 14, and 5 samples respectively.

The ecological analysis was conducted using the tissue-wide PPC. For each marker, PPC was normalized by dividing the marker’s PPC by the number of opaque pixels in the sample, thus counting only pixels that are within the tissue. The number of opaque pixels was determined using Ki67 images, as all samples had Ki67 images, and also because staining for Ki67 was conducted by one individual (A.B.). To avoid having replicates for only some samples, PPC in the two adenoma regions within each CRA and CIA sample were summed and then divided by the total number of opaque pixels in the two regions.

A distance matrix, which compares tumors based on the whole collection of markers, is required for many of the ecological tests we performed. We tested all combinations of distance metrics and normalization methods in the *vegan* R package (Oksanen et al., 2018) and found that the Jaccard distance on log-scaled PPC most often had adjacent adenomas in CIA samples as being the most similar to one another, as would be expected. It should be noted that the Jaccard distance in the *vegan* package is not the same as a traditional Jaccard distance, but more of a variant of the Bray-Curtis distance. Thus, we performed the analysis on log-scaled data and a distance matrix constructed using the Jaccard distance. PERMANOVA, PERMDSIP2, CAP, and the Mantel tests were all conducted on this distance matrix.

Spatial interaction networks were calculated using the samples that had successful alignments. In each of these samples, the PPC in each quadrat were used. Interaction networks were constructed for each sample, using two methods. In both cases, a “site” is considered to be a quadrat from the composite image. The first uses partial correlation with a spatially explicit null model. To account for spatial autocorrelation, the null model was generated using LOESS regression of the observed abundances, as suggested in (Blonder & Morueta-Holme, 2017). The R package *netassoc* was used to estimate the interaction coefficient based on partial correlation (Blonder & Morueta-Holme, 2017).

A suite of statistical tests were used to detect significant changes in marker positivity, co-localization, and direct cell-cell interactions. Several methods were used to find those results that were robust, as those that were found to be significant across multiple tests are most likely to be true. The two-sided Kruskal-Wallis rank sum test, two-sided approximate Kruskal-Wallis Test, and two-sided General Independence Test were used to determine there were differences between the tumor subtypes CRA, A-CIA, C-CIA, and CRC. Assuming that the carcinomas will take over the adenomas in the ca-in-ad samples, and thus develop into carcinomas, one can test for trends in the data by setting the timing order of A-CIA-> C-CIA - > CRC. Having ordered the subtypes, one-sided Approximative Linear-by-Linear Association Test and the Jonckheere-Terpstra test were used to determine if there a significant increase or decrease across groups (Seshan, 2018). The Conover-Iman test of multiple comparisons using rank sums was used as the post-hoc test to determine which subtypes were significantly different from one another (Dinno, 2017).

The paired-tests looking for differences between A-CIA and C-CIA included the General Symmetry test and the Wilcoxon rank sum test. As with the tests comparing all subtypes, the two-sided version of these tests was used to detect differences, and the one-sided test was used to detect the presence of trends and the direction of those trends.

The two-sided approximate Kruskal-Wallis Test, General Independence Test, General Symmetry test, and Approximative Linear-by-Linear Association Tests are permutation tests and were conducted using the *coin* package for R (Hothorn, Hornik, Wiel, & Zeileis, 2008).

### Mathematical model

A non-spatial, branching process, agent based model (ABM) was developed to simulate tumor evolution under immune predation and escape. Each agent in the model is a clone composed of cells carrying the same set of mutations. In the model each clone j from one of the three *S* _*i*_ species is composed of *N* _*ij*_ cells, where the species are: normal epithelial cells, *E*, when *i* = 0; adenoma cells, *CRA*, when *i* = 1; and carcinoma cells, *CRC*, when *i* = 2. Each species has a species-specific growth rate, *r* _*i*_, and carrying capacity, *K* _*ii*_. New clones are created during division, and, in addition to inheriting all ancestral antigens and mutations, each new clone also acquires a unique neoantigen, *γ* _*i*_, that stimulates a predatory T-cell response. Clones may protect themselves from immune attack by evolving the ability to express mechanisms that protect from immune attack, such as checkpoint inhibitors like PD-L1 or CTLA-4. Another escape route is evolving the ability to recruit immunosuppressive cells, such as neutrophils and macrophages, that not only protect from T-cell attack, but also release growth factors that can increase growth rates. New clones may also acquire driver mutations that allow the clone to evolve into a new species, with progression from *E* (0-1 driver mutations), to *CRA* (2-3 driver mutations), to *CRC* (4+ driver mutations).

Simulations were initiated with a genetically homogeneous epithelium composed of 10^7^cells. Cells died at rate of 0.01 cells per day, but divided at different rates depending on which species the clone belonged: E clones had an inherent division rate of 0.07, CRA 0.11, and CRC 0.24. The maximum size (i.e. carrying capacity) of each species also varied: the maximum number of *E* cells was 10^7^cells, 10^8^ cells for CRA, and 10^9^ cells for CRC.

In keeping with the observation that there are few driver loci in colorectal cancer [Vogelstein], the probability of “hitting” a driver gene was 0.009, meaning that the total chances of creating a clone with a new driver mutation is 2e-5. The probability of acquiring the ability to mitigate immune attack is even lower, being 0.0005, and such that the probability of creating a new clone with the ability to either suppress or block immune attack is 1e-6. New clones only acquire one new mutation, but over time it is possible to have a mutant that has accumulated all driver mutations and both escape mechanisms. Finally, it should be noted that neutral passenger mutations are not modeled, and as such, model predictions regarding genetics focus exclusively on antigenic intra-tumor heterogeneity (aITH).

The cellular mutation rate of was calculated as follows. Assuming that the exome has 2e7bp and that the base pair mutation rate per replication is 10e-10 (Lek et al., 2016; Wielgoss et al., 2011), the probability that there is at least one mutation in the exome per cell per division is 0.0027 = 1 -*Binomial k* = 0, *N* = 2*e*7, *p* = 10*e* – 10.

#### Clonal dynamics

Clonal dynamics are modeled by modifying the Lotka-Volterra competition model to create a difference equation that simulates inter- and intra-specific competition and immune predation, under the presence of genetic drift, at each time point *t* (Equation 1):

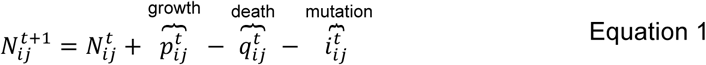

Assuming that clonal growth is logistic, the total number of new cells a clone is expected to produce during a time step is determined using Equation 2. The immunosuppressive cells recruited by clones (assuming they have evolved the ability and *e* _*ij*_ = 1) are modeled to produce growth factors, that increase that clone’s growth rate by factor ω. Random variation in the number of new cells is simulated by drawing a random variate from the Poisson distribution, using 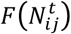 as the expected number of new cells to be created during time step *t* (Equation 3).

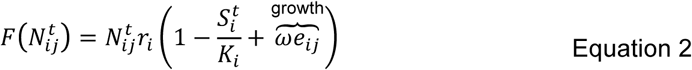

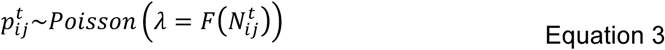

Clones lose cells in three ways. First, cells die at a constant death rate, δ. Second, clones lose cells due to competition with other species. As adenomas grow on top of the epithelium, we assume that competition between *E* and *A* is negligible. In contrast, carcinomas invade adenomas and the epithelium, and so we model competition between E & CRC, and CRA & CRC. Simulating these interactions is accomplished using the Lotka-Volterra competition model, sing the competition matrix, α (Equation 5). As 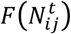 yields the number of cells that would be created in the absence of competition, and 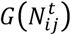 is the number of cells created in the presence of competition, 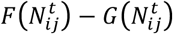 is the number of cells that were lost to competition.

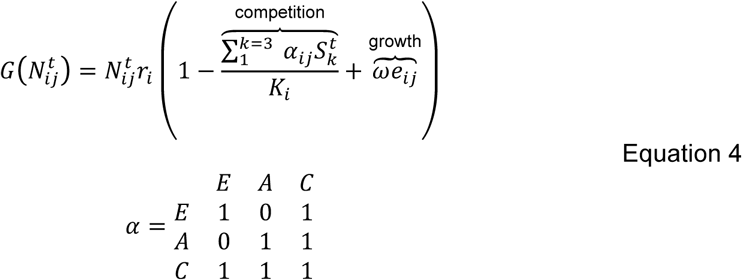

The third way in which cells may be lost is through immune predation, which removes cells at a rate proportional to the clone’s antigenicity, which is determined by the recognition potential of the clone’s neoantigen and all inherited ancestral antigens (Equation 3). However, the kill rate due to immune attack is reduced if the clone has evolved the ability to escape immune attack, either by protecting itself from T cells (i.e *d* _*ij*_ = 1), or by recruiting immunosuppressive cells (i.e. *e* _*ij*_ = 1).

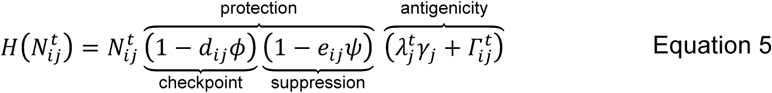

Given the three ways in which clones may be lost, the number of cells expected to die each time step is the sum of the number lost to homeostatic death, 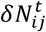, the number removed by immune predation, 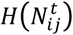, and the number lost to competition,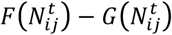. The expected number of dead cells can be used to draw a random variate from the Poisson distribution, thus adding variation in the number of cells removed due to death.

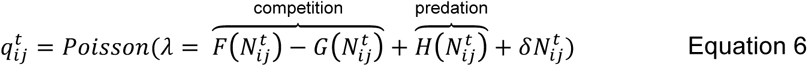

Adding random variation to the number of cells created, 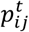, and random variation to the number lost to death, 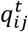, allows for the simulation of genetic drift, an important evolutionary force that plays a crucial role in determining whether or not new clones survive.

Based on the number of number of new cells created during division, 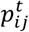, and the gene mutation rate, *μ*, clones generate 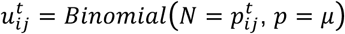 mutants each time step. Each of the 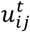 newly created clones inherits all ancestral mutations and antigens. This branching process generates a non-binary tree *T(V, E)*, in which each vertex *V*(*T*) is one of the *N* _*ij*_ clones, with edges *E*(*T*) between parental clones and each new child clone. Given this tree, each clone can access any of the ancestral clones in *ancestors*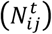, and any of the descendant clones in *descendants*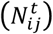. As new clones are constantly being created and removed, *ancestors*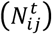, and *descendants* 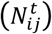 may change each time step.

#### Immune dynamics

The dynamics of T cells and immunosuppressive cells are modeled implicitly in equations 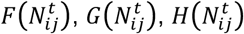 Whenever clones are created, they inherit the antigens from each of the *a*^th^ ancestors in *ancestors*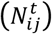, and so the clone’s initial antigenicity is 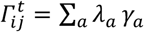. In addition to the ancestral antigens, clones also acquire a neoantigen with a random recognition potential, *γ* _*j*_ ∼*Uniform* 0,1.

We assume that there is a time lag between the clone’s time of origin, and the time at which its neoantigen is detected by the immune system. Thus, when the clone is created,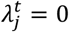. The length of the time lag is proportional to *γ*_*j*_ and the number of cells that have inherited 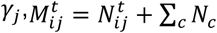 for each *c*^th^ clone in *descendants*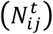, which may or may not belong to the same species as *N* _*ij*_. The probability of at least one cell carrying *γ* _*j*_ being detected is 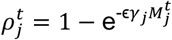, and the time *t* at which the antigen is detected, τ_*j*_, occurs when 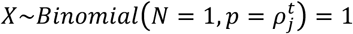. The value τ_*i*_ thus reflects the time at which the T-cell response is activated. At the time of T-cell activation, each *c*^th^ clone in *descendants* 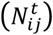 sets 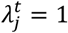 for the remainder of the simulation, and from this point on 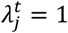 in all future descendants. In other words,

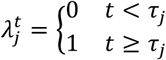

With such a time-lag, 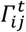 can change from one time step to another as ancestral antigens are detected, but the clone’s total antigenicity will only be to increase.

Given that T cells recognize one particular antigen, and each clone carries multiple antigens (its own neoantigen and all ancestral antigens), the total number of T cells attracted to the clone is proportional to 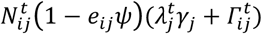. Note that clones capable of recruiting of immunosuppressive cells, i.e. those with *e*_*ij*_ = 1, are able to decreases the number of T cells attacking it. The amount of immunosuppression is determined by *ψ*, and is the same for all clones with *e*_*ij*_ = 1.

Similar to the case with T cells, the dynamics of immunosuppressive cells are assumed to follow those of the clones which are able recruit them. In this case, the number of immunosuppressive cells recruited by a clone is proportional to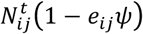.

#### Phenotypic Effects of Mutation

In addition to generating a neoantigen, *γ* _*j*_∼*Uniform* 0,1, new clones may also acquire one of three types of beneficial mutations: driver mutations, the ability to protect from T-cell attack, and the ability to recruit immunosuppressive cells. The multinomial distribution is a generalization of the binomial distribution, and is here used to determine how many of each mutant to make, including those that do not acquire beneficial mutations. Therefore *m* 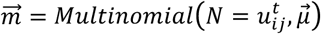 is a four element vector where: 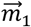 is the number of clones to create that do not have a beneficial mutation; 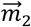 is the number of clones to create that acquire a driver mutation, with *m*_*ij*_ being the number of driver mutations *N*_*ij*_ has accumulated; 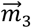 is the number of clones to create that express checkpoint inhibitors; and 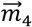 is the number of clones to create that recruit immunosuppressive cells. The parameter *n* is the number of genes in the genome, and *n*_driver_ is the number of possible driver mutations.

The vector 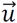 contains the probability for each mutation type, with

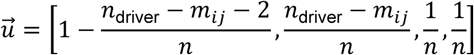

Note that calculation of 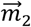 assumes the number of driver mutations that will have an effect decreases as driver mutations are accumulated. In other words, it is assumed that if a driver is “hit” more than once, the benefit comes only with the first mutation, and subsequent hits have no effect. It is also assumed there is one gene for protecting from T-cell attack, and one gene for recruiting immunosuppressive cells.

Before the creation of new clones, the number of clones created by *N*_*ij*_ at timestep *t* is 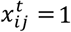. For each of the new 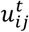 clones, an edge in the clone tree *T*(*V, E*) is created between the parent and child, *E*(*N*_*ij*_, *N*_*k,j*+x_). For new clones that do not acquire a beneficial mutation, there are no changes from the parental genotype aside from the new clone’s neoantigen. More formally, for each of the 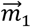 clones to be created, *d*_*k,j*+x_ = *d*_*ij*_, *e*_*k,j*+x_ = *e*_*ij*_, and *m*_*k,j*+x_ = *m*_*ij*_. Each of the 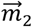 clones, which had a mutation that hit a new driver gene, *d*_*k,j*+x_ = *d*_*ij*_, and *e*_*k,j*+x_ = *e*_*ij*_, and *m*_*k,j*+x_ = *m*_*ij*_ + 1. For each 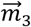 and 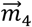 new clones to be created, *d*_*k,j*+x_ = 1, *e*_*k,j*+x_ = 1,, respectively, while *m*_*k,j*+x_ = *m*_*ij*_, for each of the 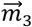 and 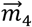 new clones. After each clone is created,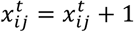. The result of this is that, after each clone is created, the total number of clones that have ever existed, regardless of species, is equal to *i*.

For the species to which the 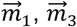 and 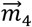 new clones belong is the same as that of the parent, i.e. *k* = *i*. However, the species *k* to which each of the 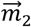 new clones belong is determined by the number of driver mutations it has accumulated:

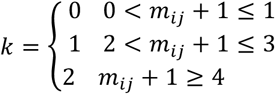

#### Simulated VAF and antigen burden calculation

In the samples, there are four different tissue types examined: adenoma (CRA), adenoma in ca-in-ad (A-CIA), carcinoma in the ca-in-ad (C-CIA), and carcinomas (CRC). Again, to be able to compare the simulated data to observed data it is necessary to look at the variant allele frequencies (VAF) in each tissue type, defined by the species they are composed of. This is straight forward in the case of CRA and CRC. However, the amount of CRA and CRC species defining CIA needs to be defined. Here, the simulated tumor is considered a CIA when *S*_1_ ≥ 1000 and *S*_2_ ≥ 1000. A-CIA is then the *S*_1_ are adenoma cells in the CIA and *S*_2_ are the C-CIA cells in the CIA. Using these definitions, each simulated tumor type, composed of species *i*, can be compared to the observed tumors of the same type.

Given the clone tree, *T*(*V, E*), it is possible to determine each antigen’s VAF, which can be compared to observed sequence data. At each time point *t*, the true VAF of each antigen *j*, which originated with clone *N*_*ij*_, in tissue type composed of species *S*_*k*_ is 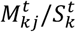, where

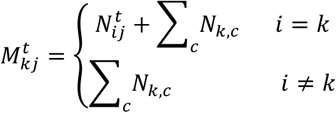

for each *c*^th^ clone in *descendants*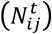.

When comparing simulated and observed VAFs, it is important to take into account of sequencing error, which here is done using the method described in (Williams, Werner, Barnes, Graham, & Sottoriva, 2016). Briefly, if the target sequencing depth is *y*, then the actual sequencing depth after accounting for error is 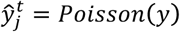. Given this sequencing depth, ŷ, the number of times each antigen is actually detected can be determined by using the binomial distribution, where the number trials is the sequencing depth, and the probability of being detected is the frequency of the mutation in the sample. Thus, the simulated VAF at time point *t* in tissue type composed of species *k* after taking sequencing error into account is 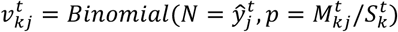

In the simulation, each antigen has a VAF for every time point it existed in a particular tissue, and so for each antigen there is a vector of VAFs over time, 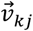. VAFs in the CRC are from the final time point, the time of resection. VAFs for CRA and CIA are determined by selecting a random day during the tumor stage.

Antigen burden in each tissue type composed of species *S*_*k*_, *B*_*k*_, can be determined using the collection of 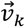 for each antigen by counting how many antigens were detected, i.e. have a VAF greater than 0. That is,

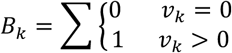

for each *v* in 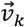.

**Supplementary table 1.** Sample details

| **Sample number** | **Classification** | **Site** | **Used for WES** | **Used for IHC** |
| --- | --- | --- | --- | --- |
| **1** | Adenoma | Oxford |  | Y |
| **2** | Adenoma | Oxford |  | Y |
| **4** | Adenoma | Oxford | Y | Y |
| **6** | Adenoma | Oxford | Y |  |
| **7** | Adenoma | Oxford |  | Y |
| **8** | Adenoma | Oxford |  | Y |
| **12** | Adenoma | Oxford | Y | Y |
| **17** | Adenoma | Oxford |  | Y |
| **22** | Adenoma | Oxford | Y | Y |
| **23** | Adenoma | Oxford |  | Y |
| **25** | Adenoma | Oxford |  | Y |
| **52** | Adenoma | UCH |  | Y |
| **57-2** | Adenoma | UCH |  | Y |
| **K** | Adenoma |  | Y |  |
| **S** | Adenoma |  | Y |  |
| **P** | Adenoma |  | Y |  |
| **31** | Ca-in-ad | Oxford |  | Y |
| **32** | Ca-in-ad | Oxford | Y | Y |
| **33** | Ca-in-ad | Oxford | Y | Y |
| **34** | Ca-in-ad | Oxford |  | Y |
| **35** | Ca-in-ad | Oxford |  | Y |
| **37** | Ca-in-ad | Oxford |  | Y |
| **38** | Ca-in-ad | Oxford |  | Y |
| **43** | Ca-in-ad | Oxford |  | Y |
| **44** | Ca-in-ad | Oxford |  | Y |
| **47** | Ca-in-ad | Oxford | Y | Y |
| **53** | Ca-in-ad | UCH |  | Y |
| **55** | Ca-in-ad | UCH |  | Y |
| **56** | Ca-in-ad | UCH |  | Y |
| **57-1** | Ca-in-ad | UCH |  | Y |
| **58-2** | Ca-in-ad | UCH |  | Y |
| **58-3** | Ca-in-ad | UCH |  | Y |
| **60** | Ca-in-ad | UCH |  | Y |
| **61** | Ca-in-ad | UCH |  | Y |
| **62** | Ca-in-ad | UCH |  | Y |
| **63** | Ca-in-ad | UCH |  | Y |
| **64** | Ca-in-ad | UCH |  | Y |
| **65** | Ca-in-ad | UCH |  | Y |
| **66** | Ca-in-ad | UCH |  | Y |
| **67** | Ca-in-ad | UCH |  | Y |
| **15** | Carcinoma | Oxford | Y | Y |
| **16** | Carcinoma | Oxford | Y | Y |
| **17** | Carcinoma | Oxford | Y | Y |
| **54** | Carcinoma | UCH |  | Y |
| **58-1** | Carcinoma | UCH |  | Y |
| **95** | Carcinoma | UCH |  | Y |
| **96** | Carcinoma | UCH |  | Y |
| **97** | Carcinoma | UCH |  | Y |
| **98** | Carcinoma | UCH |  | Y |
| **99** | Carcinoma | UCH |  | Y |
| **100** | Carcinoma | UCH |  | Y |
| **101** | Carcinoma | UCH |  | Y |
| **102** | Carcinoma | UCH |  | Y |
| **103** | Carcinoma | UCH |  | Y |
| **104** | Carcinoma | UCH |  | Y |
| **105** | Carcinoma | UCH |  | Y |
| **106** | Carcinoma | UCH |  | Y |
| **M** | Carcinoma |  | Y |  |
| **N** | Carcinoma |  | Y |  |
| **T** | Carcinoma |  | Y |  |
| **G** | Carcinoma |  | Y |  |
| **W** | Carcinoma |  | Y |  |

**Supplementary table 2.**
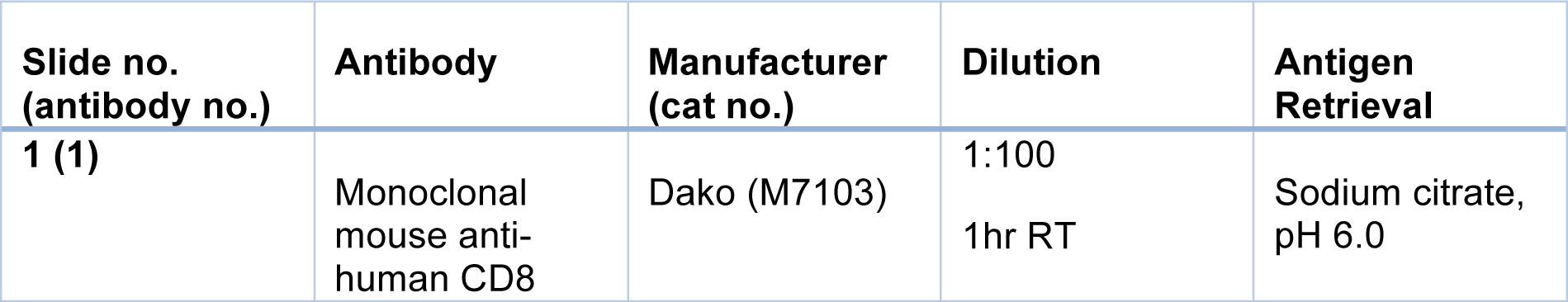
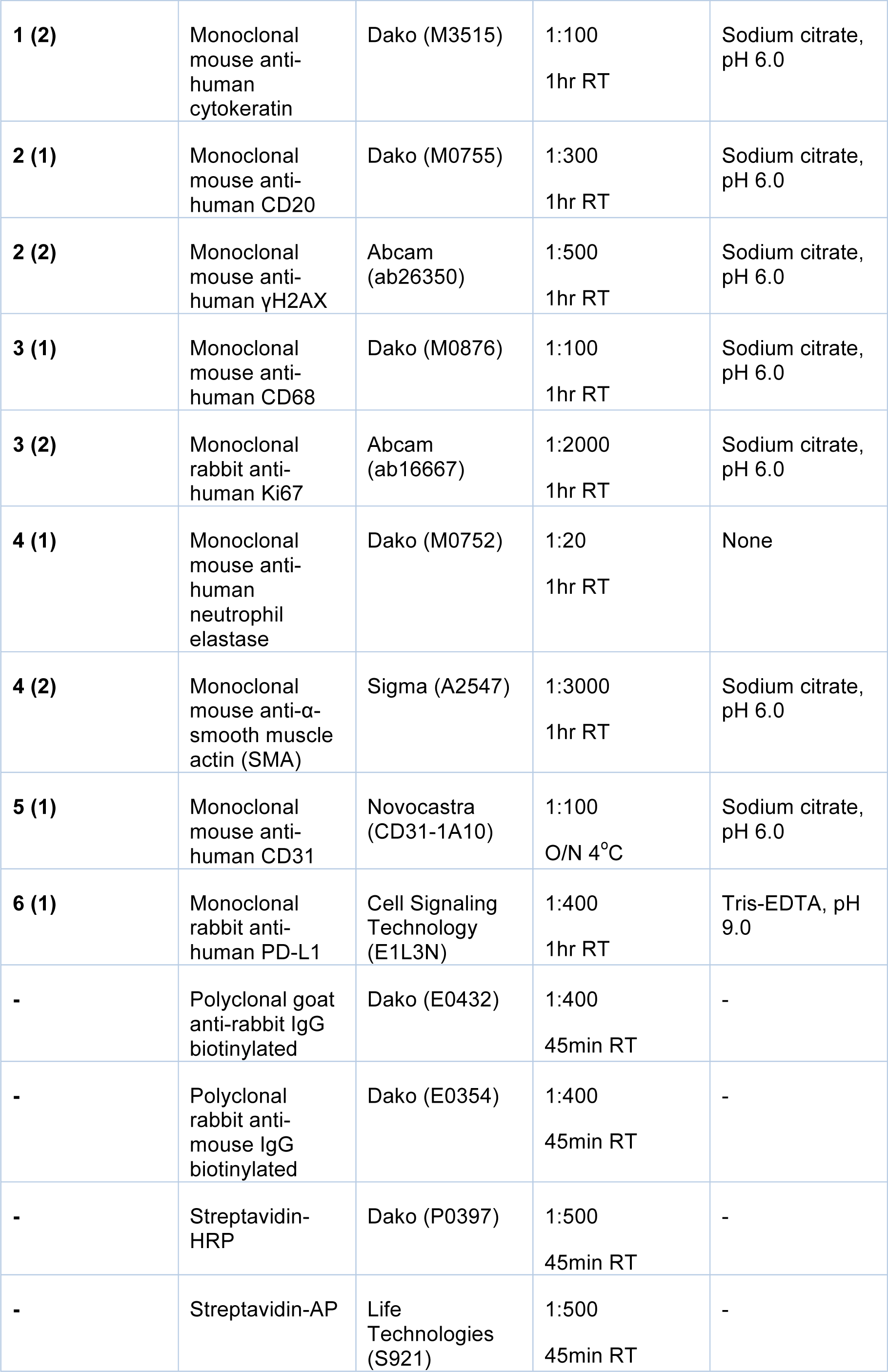
Antibody details

